# PUMA: A Phenotypic Unsupervised Model of Aging Reveals Distinct Aging Dimensions

**DOI:** 10.64898/2026.07.22.740035

**Authors:** Fatemeh Ghorbani, Ellen A.A. Nollen, Victor Guryev

## Abstract

Aging is a multidimensional process, yet most aging models reduce it to a single score to estimate biological aging and predict health outcomes, disease risk, or mortality. Here, we introduce PUMA (Phenotypic Unsupervised Model of Aging), a framework that characterizes aging through multiple phenotypic dimensions. Applying PUMA to more than 1,000 traits spanning behavioral, psychological, social, physical, environmental, and biomedical domains in over 150,000 individuals from the Lifelines cohort, we identified seven distinct phenotypic aging dimensions. These dimensions were significantly associated with the future incidence of major age-related diseases—including cancer, diabetes, COPD, heart failure, stroke, and Parkinson’s disease— demonstrating the potential of PUMA to stratify individuals by disease risk and identify phenotypic domains for targeted intervention. Notably, dimensions reflecting psychosocial factors, particularly cumulative life stress, predicted disease risk as strongly as, or more strongly than, traditional biomedical risk factors, highlighting the importance of psychological and social influences in aging and disease risk. The observation that phenotypic aging dimensions are differentially associated with the future risk of age-related diseases supports a multidimensional model of aging, indicating that aging is not a single uniform process. Because PUMA relies on accessible phenotypic data, it provides an interpretable framework for disease risk stratification and targeted preventive strategies.

## Introduction

Aging is increasingly recognized as a complex, multifactorial, and dynamic process, shaped by interactions among biological, psychological, behavioral, social, and environmental factors^1-4^. This recognition has led to growing interest in more integrative perspectives, ones that view aging not as the result of a single cause, but as the outcome of many interacting processes over time^5^. Reflecting this shift, numerous large-scale longitudinal cohorts have been established, enabling the study of aging across multiple domains^6^.

Yet despite this recognition and the availability of rich data, many studies continue to adopt reductionist approaches, focusing on isolated factors and missing the complex patterns that characterize the aging process^1, 7^. Thus, approaches that account for the complexity of aging are needed to complement existing models, offering a more integrated view of how diverse processes interact over time. Most studies aiming to capture the complexity of aging have relied on molecular data, such as epigenetic, proteomic and metabolomic data to estimate overall biological age^8-11^. Far fewer have extended this approach to phenotypic data, particularly clinical, physiological, or physical functioning measures^12-14^. Among those that do, most rely on predefined or narrowly selected phenotypic variables, which limits their ability to capture the full complexity and heterogeneity of aging across domains. Furthermore, most existing models— whether biological or phenotypic—tend to compress these inputs into a single score, reducing aging to a unidimensional measure and overlooking its multidimensional nature.

To address these limitations, we introduce PUMA (Phenotypic Unsupervised Model of Aging), a novel, unsupervised approach that uses rich phenotypic data from the Lifelines cohort, one of the most phenotypically comprehensive population-based studies worldwide. Unlike existing models that rely on predefined phenotypic domains or reduce aging to a single composite score, PUMA identifies multiple phenotypic aging dimensions derived from more than a thousand behavioral, psychological, social, physical, environmental, and biomedical variables.

Each dimension captures a distinct pattern of age-associated phenotypic variation, providing a multidimensional view of aging. We further demonstrate the clinical relevance of these dimensions through their significant associations with the future onset of major age-related diseases. These associations not only enable disease risk stratification but also identify the phenotypic domains most strongly associated with each condition. PUMA therefore advances a multidimensional understanding of aging while providing a practical framework for disease risk stratification and targeted prevention strategies.

## Methods

### Study cohort

We used data from the Lifelines Cohort Study, a multidisciplinary, population-based prospective cohort comprising 167,729 individuals with ongoing longitudinal follow-up^15^. Participants were enrolled between 2006 and 2013 (baseline assessment), with follow-up assessments conducted in 2014–2017 (second assessment) and 2019–2023 (third assessment)^16^. Baseline data were available for 167,729 participants, for whom 2,232 phenotypic variables spanning behavioral, psychological, social, environmental, physical, and biomedical domains were collected^17^. To ensure sufficient data coverage, data-availability filters were applied to both participants and variables. Details of the filtering procedure are provided in Supplementary Note 1.

The Lifelines Cohort Study received approval from the Medical Ethics Committee of the University Medical Center Groningen. Data were accessed via the Lifelines data catalogue (https://data-catalogue.lifelines.nl; accessed January 2023; project ID OV21_00356).

### Identification of aging-associated phenotypic dimensions

A modified version of Principal Component Analysis (PCA) capable of handling missing data, known as PCA of Incomplete Data (InDaPCA)^18^, was applied to the baseline dataset. In InDaPCA, covariances are computed using the available data for each pair of variables, allowing missing values to be skipped, and the resulting eigenvalues and eigenvectors are used to calculate component scores^18^. Spearman rank correlation was used to assess the significance of the association between each principal component (PC) score and age. PCs were considered aging-associated if they explained more than 1% of the total variance and exhibited a nominally significant association with age (Spearman p < 0.05). Each aging-associated PC was interpreted as a distinct phenotypic aging dimension representing coordinated variation across multiple phenotypic traits.

### Characterization of phenotypic aging dimensions

To identify the baseline variables most strongly associated with each aging-associated PC, partial correlation analysis was performed. Partial correlations were computed using the ppcor package in R ^19^. This method quantifies the relationship between each phenotypic variable and a given PC while controlling for the effects of all other variables in the dataset. For each PC, variables were ranked based on the magnitude and statistical significance of their partial correlation coefficients. The highest-ranking variables were retained as key contributors, reflecting the strongest associations with that component. To characterize the major phenotypic features of each PC, the key variables were classified into predefined Lifelines catalogue subsections. For each PC, the subsection containing the largest number of key variables was assigned as its representative phenotypic domain. A chi-square test was used to assess whether key variables were overrepresented within a subsection relative to its overall frequency in the Lifelines catalogue. Subsections were considered enriched if they showed nominally significant overrepresentation (p < 0.05). The representative phenotypic domain formed the basis for interpreting the corresponding phenotypic aging dimension.

### Composite score calculation

For each aging-associated PC, we performed multiple linear regression using its key contributing variables as predictors and the corresponding InDaPCA-derived PC scores as the dependent variable. The resulting values represent individual-level composite scores for each component (e.g., PC1-CS, PC2-CS). To account for age-related differences and facilitate comparability across age groups, composite scores were standardized within each 10-year age bin separately using z-score normalization^20^. This procedure standardizes score distributions across age groups, allowing composite scores to reflect relative phenotypic variation within age groups while reducing the influence of age-related differences.

### Assessment of disease incidence

To evaluate the clinical relevance of phenotypic aging dimensions, we assessed incident cases of six major age-related diseases among baseline participants: cancer, diabetes, COPD, heart failure, stroke, and Parkinson’s disease. Disease incidence was determined longitudinally using self-reported questionnaires collected during the second and third Lifelines follow-up assessments. At each follow-up, participants were asked whether they had been diagnosed with any of these conditions since their previous assessment. A disease was classified as incident if it was newly reported at either follow-up among individuals without a history of that condition at baseline. The specific coding scheme used to identify incident cases is detailed in Supplementary Tables 1 and 2.

### Clinical relevance analyses

To assess whether the phenotypic aging dimensions were associated with incident disease, we applied an enrichment-based approach adapted from the Gene Set Enrichment Analysis (GSEA)^21^ framework. This method evaluates whether incident cases of a given disease are statistically overrepresented among participants with high or low scores along each phenotypic aging dimension. For this purpose, we used the previously computed composite scores (PC-CS), which quantify each participant’s position on the corresponding dimension.

Participants were ranked in descending order according to their age-adjusted PC-CS for each dimension, such that individuals with the highest scores were positioned at the beginning (left side) of the ranked list. A running sum statistic was then calculated across the ranked list, increasing when a participant with incident disease was encountered and decreasing when a participant without incident disease was encountered. The maximum deviation from zero, referred to as the enrichment score (ES), indicates whether disease incidence is concentrated among participants with high (positive ES) or low (negative ES) dimension scores. To allow comparison across dimensions and diseases, the ES was normalized to obtain a normalized enrichment score (NES), adjusting for differences in score distributions. Statistical significance of the NES was determined using permutation testing. P-values were corrected for multiple comparisons using the Benjamini–Hochberg method to control the false discovery rate (FDR), and adjusted p-values < 0.05 were considered statistically significant. For each significant dimension–disease association, the representative domain assigned during dimension characterization was used to identify the phenotypic features most strongly linked to the disease. To assess whether phenotypic aging dimensions could stratify individuals by their risk of developing diseases, we estimated relative risk (RR) for each dimension that showed a significant association with disease. High-risk groups were defined using the leading-edge subset of participants contributing most to the enrichment signal, comprising individuals with extreme composite scores at either end of the dimension score distribution where disease cases were concentrated. For dimensions with a positive enrichment score, the high-risk group comprised individuals with the highest PC-CS values; for negative enrichment scores, the group included those with the lowest scores. The low-risk group consisted of participants with intermediate composite scores, where no enrichment was observed. Disease incidence within each group was calculated as the number of individuals with incident disease divided by the total number in the group. RR was then calculated as the ratio of disease incidence in the high-risk group to disease incidence in the low-risk group.

## Results

### Phenotypic aging dimensions

To identify distinct phenotypic aging dimensions, we applied PUMA (Phenotypic Unsupervised Model of Aging), an unsupervised approach, to a large set of multidomain phenotypic variables collected at baseline in the Lifelines cohort. An overview of the study design and analytical workflow is shown in Figure 1A. After filtering participants and phenotypic variables based on data availability, 152,241 participants aged 18 years and older and 1,177 variables were retained for analysis (Figure 1B). Of these variables, 31% were directly measured, and 69% were obtained from questionnaires.

**Figure 1.**
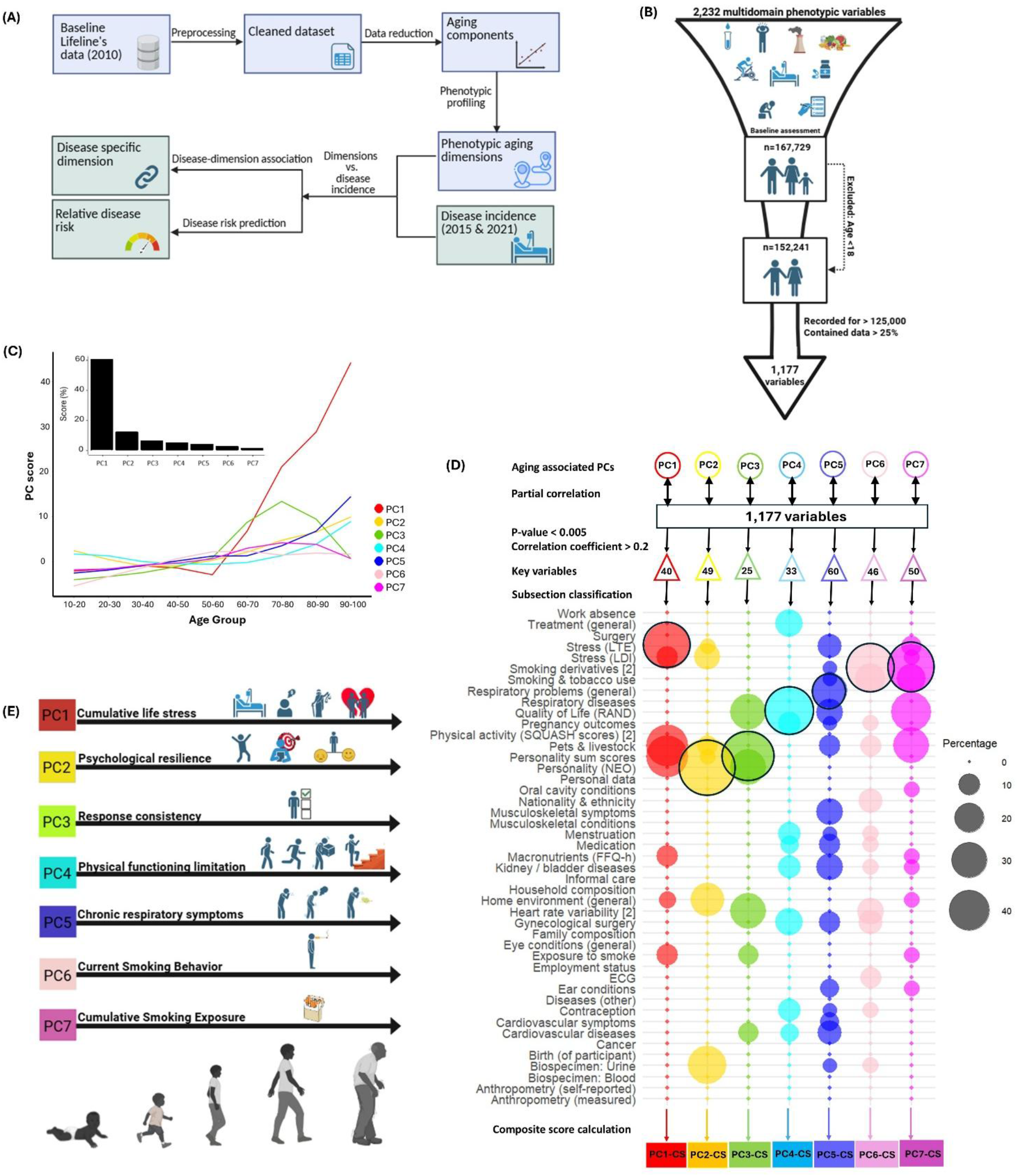
Phenotypic aging dimensions identified by PUMA. **(A)** Overview of the PUMA (Phenotypic Unsupervised Model of Aging) analytical workflow, illustrating the two main phases: extraction of phenotypic aging dimensions (blue boxes) and evaluation of their clinical relevance (green boxes). **(B)** Data filtering procedure. A total of 152,241 baseline participants aged 18 years and older were retained for analysis. In parallel, 1,177 phenotypic variables were included based on availability in more than 125,000 participants and more than 25% non-missing data. **(C)** Aging-associated PCs. PC1–PC7 each explained more than 1% of the total variance and were significantly associated with age (p-value < 0.05). The line plot depicts mean PC scores across age groups and inset bar plot shows the proportion of variance explained by each component. **(D)** Phenotypic characterization of PC1–PC7. Key variables for each component were identified using partial correlation analysis and classified into predefined Lifelines catalogue subsections. The subsection containing the largest number of key variables (black circle) was designated as the representative phenotypic domain. Composite scores were then calculated from the key variables for each PC and standardized by age to account for age-related differences. **(E)** Descriptive phenotypic labels for PC1–PC7. Key variables within the representative phenotypic domain of each aging-associated PC were reviewed to assign descriptive labels, defining distinct phenotypic aging dimensions.

Principal Component Analysis (PCA) was applied to the filtered baseline dataset to reduce dimensionality and identify major patterns of phenotypic variation across individuals. The first seven principal components (PC1–PC7) were retained based on an explained variance threshold greater than 1%. Collectively, these components accounted for over 92% of the total variance in the data, with individual contributions of 60.6% (PC1), 12.2% (PC2), 6.4% (PC3), 4.9% (PC4), 4.1% (PC5), 2.7% (PC6), and 1.3% (PC7). Additionally, all seven components were significantly associated with age (p-value < 0.05) and were therefore classified as aging-associated PCs (Figure 1C). These aging-associated PCs formed the basis for the phenotypic aging dimensions characterized in subsequent analyses. The distributions of the seven aging-associated PCs across age groups, stratified by sex, are shown in Supplementary Figure 1.

To identify the key phenotypic variables associated with each aging-associated PC, we conducted partial correlation analyses between PC1–PC7 and the 1,177 variables. Key variables were selected using data-driven thresholds (p < 0.005; partial correlation > 0.2), yielding between 25 and 60 variables per aging-associated PC. These key variables were then classified into predefined Lifelines subsections to characterize the phenotypic basis of each aging-associated PC (Figure 1D). For each PC, the subsection containing the largest number of key variables was assigned as its representative phenotypic domain. For PC1 through PC5, the representative domains were Stress (LTE), Personality (NEO), Personality Sum Score, Quality of Life (RAND), and Respiratory Problems. PC6 and PC7 were both primarily related to the Smoking Derivatives domain. Definitions of Lifelines subsections and the corresponding key variables for each PC are provided in Supplementary Tables 3 and 4.

However, the representative phenotypic domains derived from Lifelines subsections were often broad and did not fully capture the specific phenotypic characteristics reflected in each component. To improve interpretability, we reviewed the key variables within each representative subsection and assigned a descriptive label that more precisely reflected its underlying phenotypic content. These refined interpretations enabled us to define each aging-associated PC as a distinct phenotypic aging dimension (Figure 1E):

- **PC1 – Cumulative Life Stress:** Based on variables assessing exposure to major stressors such as bereavement, serious illness or injury, financial hardship, legal difficulties, and interpersonal conflict.
- **PC2 – Psychological Resilience:** Capturing personality traits related to emotional stability, optimism, and goal-directedness.
- **PC3 – Response Consistency:** Reflecting the completeness and consistency of questionnaire responses, potentially serving as an indirect indicator of engagement during data collection.
- **PC4 – Physical Functioning Limitations:** Reflecting limitations in daily physical activities, including walking, stair climbing, lifting, and self-care.
- **PC5 – Chronic Respiratory Symptoms:** Involving persistent coughing, phlegm production, and breathlessness indicative of pulmonary dysfunction.
- **PC6 – Current Smoking Behavior:** Capturing indicators of recent or ongoing tobacco use, including current smoking status, amount smoked per day, and recent initiation.
- **PC7 – Cumulative Smoking Exposure:** Reflecting long-term patterns such as smoking duration, lifetime quantity, and historical use.

To quantify the phenotypic aging dimensions at the individual level, composite scores were calculated for each baseline participant using the key variables associated with PC1–PC7. These scores (PC1-CS to PC7-CS) represent an individual’s position along each phenotypic aging dimension. To account for age-related differences, scores were standardized within age groups, allowing them to capture phenotypic variation independent of chronological age.

### Clinical relevance of the phenotypic aging dimensions

#### -Associations between phenotypic aging dimensions and disease outcomes

Having identified seven aging-associated phenotypic dimensions, we next evaluated their clinical relevance by testing associations with the incidence of major age-related diseases: cancer, diabetes, COPD, heart failure, stroke, and Parkinson’s disease. Incident disease cases were identified using self-reported data from the second and third follow-up assessments, capturing new diagnoses reported after baseline. Of the 152,241 participants assessed at baseline, approximately 113,000 attended at least one follow-up and were eligible for disease incidence analysis. The number of participants with available data for each condition, along with the count of incident cases, is summarized in Figure 2A.

**Figure 2.**
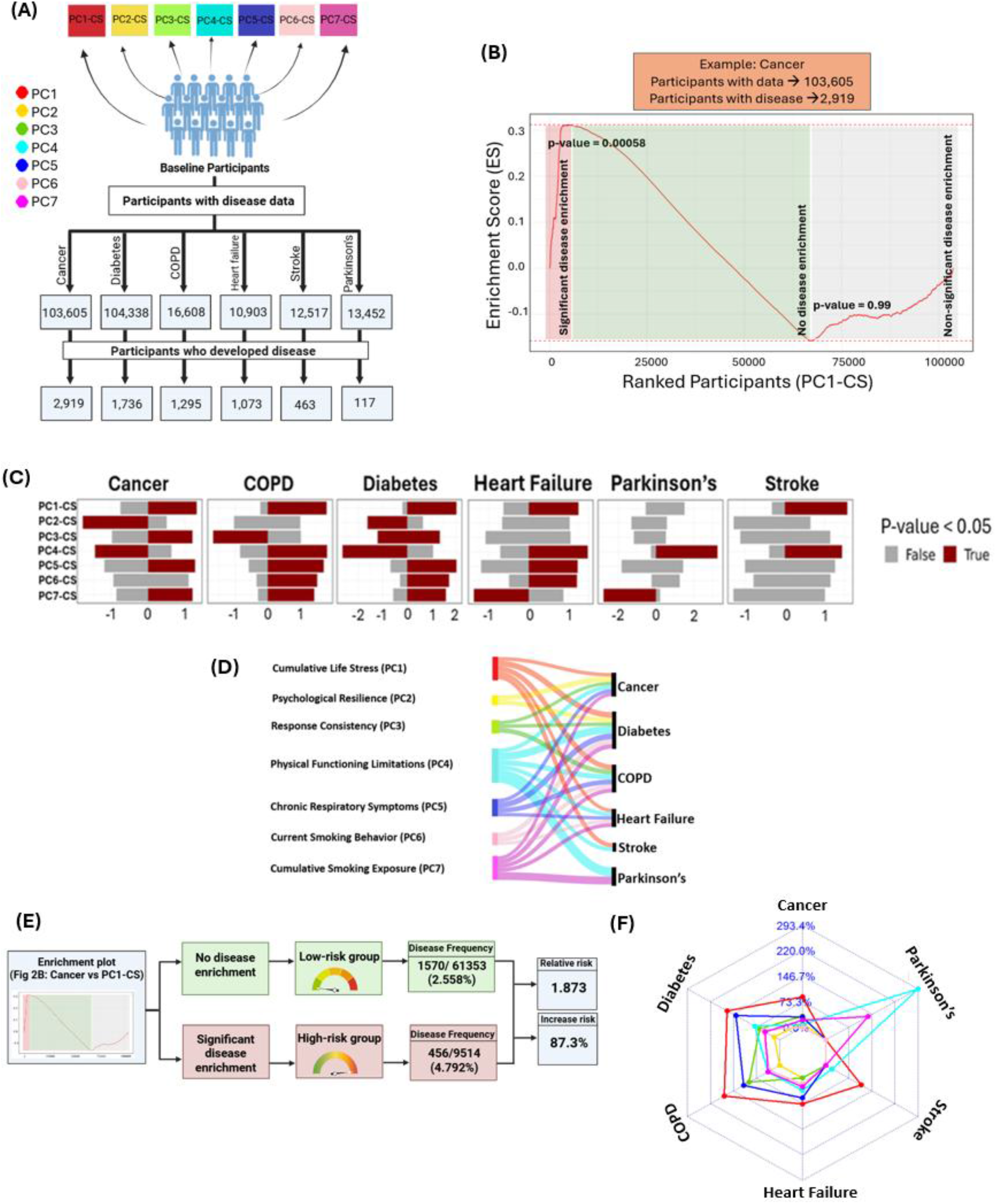
Clinical relevance of the phenotypic aging dimensions. **(A)** Overview of baseline participants with disease follow-up data and incident cases for cancer, diabetes, COPD, heart failure, stroke, and Parkinson’s disease. Composite scores (PC1–PC7-CS) were calculated for each individual to quantify their position along the phenotypic aging dimensions, enabling assessment of their associations with incident disease. **(B)** Illustrative example of enrichment analysis for cancer incidence across participants ranked by PC1 composite scores (PC1-CS). A total of 103,605 participants with cancer data are ordered from highest to lowest PC1-CS. The enrichment score (ES) reflects the maximum deviation from zero, indicating whether cancer incidence is concentrated toward the high or low end of the PC1-CS distribution. A significant positive enrichment score (p = 0.00058; red shaded region) indicates overrepresentation of cancer cases among participants with high PC1-CS values. In contrast, a non-significant negative enrichment score (p = 0.99; gray shaded region) indicates no meaningful enrichment at the lower end of the score distribution. **(C)** Summary of enrichment results across the seven phenotypic aging dimensions (PC1–PC7) for each disease. Bars represent normalized enrichment scores (NES), indicating the strength and direction of disease incidence enrichment along the composite score distribution. Positive values reflect enrichment among individuals with high scores; negative values reflect enrichment among those with low scores. Statistically significant associations (FDR-adjusted p-value < 0.05) are shown in red; non-significant associations are shown in gray. While panel B illustrates the raw ES for a single example (cancer and PC1), panel C summarizes the analysis across all phenotypic aging dimensions and diseases using NES, enabling direct comparison across dimensions and diseases. **(D)** Key phenotypic domains associated with disease incidence, derived from the descriptive labels assigned to phenotypic aging dimensions showing significant disease associations. Line width reflects the magnitude of the NES. **(E)** Illustrative example of relative risk (RR) estimation for cancer based on PC1-CS. The high-risk group comprises participants in the red shaded region that contribute most to the enrichment signal, while the low-risk group is drawn from the middle (green) region of the distribution where no enrichment is observed. Disease frequency, RR, and percentage increase in risk are calculated by comparing incidence between these two groups. **(F)** Radar plot summarizing percentage increase in disease risk across phenotypic aging dimensions. Each axis represents one disease, with radial distance indicating the percentage increase in incidence among high-risk individuals. Colored lines represent the contribution of each dimension (PC1–PC7).

To test whether the identified phenotypic aging dimensions were associated with disease incidence, we used participant-level composite scores (PC1-CS to PC7-CS), which quantify an individual’s position along each dimension. Participants were ranked according to their age-adjusted scores, and an enrichment-based approach was used to determine whether incident disease cases were overrepresented at either end of the score distribution. A running sum statistic was calculated across the ranked list, increasing when an incident case was encountered and decreasing otherwise. The maximum deviation from zero, termed the enrichment score (ES), indicates whether disease incidence was overrepresented among individuals with particularly high (positive ES) or low (negative ES) scores. An illustrative example is shown in Figure 2B, where incident cancer is significantly overrepresented among individuals with high scores on the cumulative life stress dimension (PC1-CS), resulting in a positive enrichment score. A complete summary of enrichment scores and their significance across all phenotypic aging dimensions and diseases is provided in Supplementary Table 5 and Supplementary Figure 2.

To enable comparison across dimensions and diseases, raw enrichment scores were normalized to produce normalized enrichment scores (NES), which reflect both the strength and direction of association while accounting for differences in score distributions. Phenotypic aging dimension scores showed significant associations with the future onset of age-related diseases detected during follow-up, demonstrating their clinical relevance. These dimension–disease associations are summarized in Figure 2C.

#### -Key phenotypic domains associated with disease

Building on the significant dimension–disease associations (Figure 2C), we next identified the phenotypic domains most strongly associated with each disease using the descriptive labels assigned to each phenotypic aging dimension (Figure 2D). Each disease was associated with a distinct combination of phenotypic domains. Cancer incidence was associated with cumulative life stress, psychological resilience, response consistency, physical functioning limitations, and chronic respiratory symptoms. COPD was associated with cumulative life stress, chronic respiratory symptoms, physical functioning limitations, current smoking behavior, and cumulative smoking exposure. Diabetes showed similarly broad associations spanning cumulative life stress, psychological resilience, physical functioning limitations, chronic respiratory symptoms, and smoking-related domains. Heart failure was primarily associated with cumulative life stress and physical functioning limitations, with additional associations with chronic respiratory symptoms and cumulative smoking exposure. Stroke was associated mainly with cumulative life stress and physical functioning limitations, whereas Parkinson’s disease was associated with physical functioning limitations and cumulative smoking exposure.

#### -Stratification of individuals by relative disease risk

To evaluate the predictive utility of the phenotypic aging dimensions, we assessed whether they could stratify individuals according to their relative risk (RR) of developing disease. For each significant dimension–disease association (Figure 2C), RR was estimated by comparing disease incidence between high- and low-risk groups defined by participants’ positions along the composite score distribution. We also calculated the corresponding percentage increase in risk to quantify the magnitude of the difference between the two groups. An illustrative example is shown in Figure 2E, where individuals with high scores on the cumulative life stress dimension (PC1-CS) had an approximately 90% higher risk of developing cancer than those in the low-risk group. A complete overview of risk estimates for all significant dimension–disease associations is provided in Supplementary Table 6, with a summary of the relative risk increases shown in Figure 2F.

The magnitude of the relative risk increase varied considerably across phenotypic aging dimensions and diseases (Figure 2F). The cumulative life stress dimension (PC1) showed the largest increases in risk across multiple diseases, including cancer, diabetes, COPD, heart failure, and stroke. In contrast, the physical functioning limitations dimension (PC4) was associated with the greatest increase in risk for Parkinson’s disease, highlighting the importance of impaired physical functioning in this condition.

## Discussion

In this study, we developed PUMA (Phenotypic Unsupervised Model of Aging), an unsupervised framework to characterize the multidimensional nature of human aging. Using more than 1,000 phenotypic variables spanning behavioral, psychological, social, physical, environmental, and biomedical domains in over 150,000 individuals, we identified seven phenotypic aging dimensions. Rather than reducing aging to a single composite score, these dimensions capture distinct aspects of phenotypic aging that vary across individuals. Importantly, each dimension was significantly associated with the future incidence of major age-related diseases, including cancer, diabetes, COPD, heart failure, stroke, and Parkinson’s disease. These findings demonstrate the clinical relevance of the identified phenotypic aging dimensions for disease risk stratification and highlight distinct phenotypic domains that may inform targeted prevention strategies. Overall, PUMA provides an unsupervised framework for modeling phenotypic aging that captures both the multidimensional nature of aging and its clinical relevance.

Aging clocks are commonly categorized as molecular or phenotypic. Molecular clocks, based on data such as DNA methylation^8, 9, 27-29^, proteomics^10, 25, 26^, or metabolomics^11, 22-24^ are powerful but require specialized assays. In contrast, phenotypic clocks typically rely on routinely collected, non-invasive, and interpretable clinical and physiological data^12-14, 30-36^. These models not only outperform chronological age in predicting mortality but are also more sensitive to behavioral and lifestyle changes, making them more actionable and scalable for clinical practice^37^. Despite these advantages, phenotypic clocks remain limited in scope. Most existing models rely on a set of predefined clinical and physiological indicators such as blood biomarkers, blood pressure, and body mass index (BMI), while overlooking other domains of phenotypic aging. In addition, many approaches reduce aging to a single composite score, which, although practical, may obscure the multidimensional and heterogeneous nature of aging across individuals and phenotypic domains. PUMA expands the scope of phenotypic aging research by applying an unsupervised approach to more than 1,000 phenotypic traits. Unlike previous models that focus primarily on physiological indicators, PUMA integrates a much broader range of traits, including behavioral, psychological, social, physical, environmental, and biomedical that, to our knowledge, have not previously been incorporated into a single phenotypic aging framework. Furthermore, PUMA adopts an explicitly multidimensional approach, identifying multiple independent phenotypic aging dimensions that each capture a distinct aspect of aging. This enables a more interpretable representation of aging by capturing heterogeneity both within and between individuals. Only a few previous phenotypic aging studies have adopted a similarly multidimensional perspective, including those by Farrell et al.^13^ and Salimi et al.^12^. However, PUMA differs in both scope and purpose by using a substantially broader range of phenotypic traits and linking distinct aging dimensions to future disease risk.

A key finding of this study is the strong clinical relevance of phenotypic aging dimensions representing non-biomedical domains, including cumulative life stress, psychological resilience, and response consistency, for major age-related diseases. Although these domains are rarely incorporated into phenotypic aging models, they showed disease associations comparable to, and in some cases stronger than, those of traditional biomedical domains, including smoking, respiratory symptoms, and physical functioning limitations. Their identification highlights the importance of underexplored psychosocial and behavioral aspects of aging and suggests new opportunities for disease prevention beyond conventional biomedical risk factors.

Among the seven phenotypic aging dimensions, PC4, representing physical functioning limitations, was significantly associated with all six diseases, consistent with its well-established relationship with morbidity and mortality in aging populations. PC1 and PC7, representing cumulative life stress and cumulative smoking exposure, respectively, were each associated with five of the six diseases. While smoking is a well-established risk factor for multiple chronic diseases, the contribution of cumulative life stress to disease risk has received comparatively less attention. Notably, the cumulative life stress dimension showed the largest increases in relative risk across five diseases, suggesting that stress-related factors contribute substantially to inter-individual differences in aging. Biologically, chronic stress affects immune^38^, metabolic^39^, and inflammatory pathways^40^, contributing to increased vulnerability to chronic disease. These findings reinforce growing evidence linking life stress to disease risk and underscore the potential value of stress-reduction strategies for healthy aging and disease prevention.

The evaluation of sex-specific differences in relative disease risk showed that, for most phenotypic aging dimensions, the magnitude of risk stratification was similar between females and males across all diseases. However, PC4 consistently showed higher relative risks in females than in males across all six diseases, particularly for cancer and diabetes, for which no significant increase in relative risk was observed in males. This difference may reflect the composition of PC4, which included a substantial number of variables related to female-specific characteristics, such as pregnancy outcomes, menstruation, gynecological surgeries, and contraceptive use. Sex-specific relative risk estimates are provided in Supplementary Table 6.

A particularly intriguing finding of this study was PC3, the response consistency dimension, which was characterized by patterns in participants’ questionnaire responses rather than a conventional phenotypic domain. Whether this dimension reflects participant engagement, response behavior, cognitive function, or other methodological or biological factors remains unclear and warrants further investigation.

This study also highlights that phenotypic aging dimensions differ in their associations with specific diseases. For example, cancer risk was most strongly associated with cumulative life stress, whereas Parkinson’s disease was most strongly associated with physical functioning limitations. These findings support the concept that aging is not a single, uniform process but rather comprises multiple phenotypic dimensions that contribute differently to disease risk. Unlike traditional approaches that reduce aging to a single composite score, PUMA captures this multidimensionality by identifying distinct phenotypic aging dimensions, providing a more comprehensive framework for understanding disease risk.

Importantly, although each phenotypic aging dimension was assigned a descriptive label based on its dominant phenotypic feature (e.g., cumulative life stress, psychological resilience, or physical functioning limitations), each dimension represents a multivariate pattern rather than an isolated trait. Consequently, high or low scores on a given dimension reflect the combined influence of multiple correlated variables. For example, the dimension labeled “physical functioning limitations” showed a negative association with cancer and diabetes. While this may appear unexpected if interpreted solely in terms of physical functioning, the dimension also captures additional correlated variables that contribute to its overall composition. Therefore, the observed associations should be interpreted as reflecting the phenotypic aging dimension as a whole rather than the effect of any single variable. These findings underscore the importance of interpreting dimension–disease associations as multivariate phenotypic patterns rather than direct effects of individual traits.

There is substantial evidence that socioeconomic status (SES), typically measured through education and wealth, influences aging^41^. Consistent with this, PC1, PC3, PC5, and PC7 included variables related to SES, particularly indicators of employment status and financial difficulties. Similarly, growing evidence suggests that dietary factors contribute to aging and age-related health outcomes.^42^ In line with this, PC1 and PC7 also included nutrition-related variables, including dietary patterns. Although SES- and diet-related variables did not constitute the dominant phenotypic domains of these dimensions, their presence suggests that they contribute to the broader phenotypic processes captured by the identified aging dimensions rather than representing independent phenotypic aging dimensions themselves.

Theoretically, our findings support a multidimensional model of aging that contrasts with traditional unidimensional approaches. The identification of distinct phenotypic aging dimensions, each showing different associations with age-related diseases, reinforces the view that aging comprises multiple interconnected processes rather than a single, uniform process. This perspective is consistent with the Heterogeneity of Aging framework, which proposes that aging does not follow a single trajectory^43^. Furthermore, our findings align with life-span developmental theory, which views aging as a multidimensional process shaped by biological, psychological, and social influences throughout life^44^. The prominent role of dimensions related to cumulative life stress and psychological resilience further emphasizes the importance of psychological factors in aging and age-related disease risk.

The PUMA framework relies on routinely collected, low-cost, and interpretable phenotypic data, making its potential integration into clinical settings such as general practice or public health screening feasible. By collecting the phenotypic variables required for each dimension and using them as input to PUMA, dimension-specific scores can be computed and compared with risk thresholds to identify individuals at elevated disease risk. These thresholds are derived from dimension-specific composite scores at the point in the distribution where significant disease enrichment occurs and are detailed in Supplementary Table 6. Unlike conventional aging models that define ‘high risk’ based on arbitrary cutoffs (e.g., top quartile), PUMA derives dimension-specific thresholds based on disease enrichment, enabling more targeted risk stratification. High-risk individuals may be interpreted as exhibiting accelerated aging within a specific phenotypic domain, meaning that they display phenotypic patterns associated with an elevated risk of certain diseases. By identifying the phenotypic domains in which an individual exhibits the greatest phenotypic deviation, PUMA has the potential to support more personalized prevention strategies targeting the phenotypic domains most strongly associated with future disease risk. This domain-specific approach may facilitate earlier risk identification and more targeted prevention.

Before PUMA can be applied in clinical settings, its predictive validity should be evaluated in independent and diverse populations. Although developed using the Lifelines cohort, the framework may require adaptation to account for population-specific phenotypic characteristics.

Future studies should therefore validate PUMA in independent cohorts, such as the UK Biobank, to assess its generalizability and robustness across different populations. While the identified phenotypic aging dimensions may differ between populations, the broader objective is to validate the underlying framework by demonstrating that multidimensional phenotypic aging consistently predicts disease risk across diverse populations. Ultimately, the most useful implementation of PUMA will be one tailored to the characteristics of the target population.

This study has several limitations that also highlight important directions for future research. First, the phenotypic aging dimensions were derived from cross-sectional data, capturing differences between individuals rather than changes within individuals over time. Longitudinal analyses are needed to determine how these phenotypic aging dimensions evolve within individuals over time. Second, the current framework relies on linear PCA to derive phenotypic aging dimensions. While effective at capturing major sources of variation, linear PCA may not fully represent non-linear age-related patterns present in the data. Future studies should evaluate non-linear dimensionality reduction approaches to improve the representation and interpretability of phenotypic aging dimensions. Third, disease outcomes were based on self-reports during follow-up assessments. Although self-report is widely used in large-scale studies, it may be subject to bias. Future validation using clinically verified diagnoses or medical records would strengthen confidence in the observed associations. Fourth, while the Lifelines cohort provided a rich foundation for developing PUMA, validation in independent populations is necessary to establish its generalizability. Fifth, as with any observational study, the findings reported here are correlational and do not establish causality. Unmeasured confounding factors could influence both phenotypic aging and disease outcomes. For example, limited access to healthcare may accelerate phenotypic aging by increasing unmanaged stress, while also delaying disease detection. Future studies should use statistical methods designed to assess causality to better understand the mechanisms linking phenotypic aging dimensions to disease risk. Sixth, the current analysis assumes that each aging dimension contributes independently to disease risk. However, interactions between dimensions may increase vulnerability. For example, the combination of cumulative life stress and physical functioning limitations may increase cardiovascular disease risk more than either dimension alone. Exploring such interactions could reveal synergistic effects and offer deeper insights into health risks. Finally, future studies should explore the integration of phenotypic aging dimensions with molecular markers to strengthen mechanistic interpretation. Integrating genomic, epigenomic, and other molecular layers may help uncover the biological mechanisms underlying the identified phenotypic aging dimensions. In conclusion, this study demonstrates that aging is best understood not as a single, uniform process, but as a collection of distinct phenotypic dimensions that span different domains. PUMA captures this complexity by identifying multiple phenotypic aging dimensions, each associated with disease risks, providing a more precise and interpretable representation of how individuals age. Because it relies on accessible, non-invasive phenotypic data, PUMA has the potential to support disease risk stratification and domain-specific prevention. With future validation in independent cohorts and integration with molecular aging markers, PUMA provides a foundation for advancing our understanding of aging heterogeneity and informing future strategies for the prevention of age-related diseases.

## Data availability

The data used in this study were obtained from the Lifelines Cohort Study under project ID OV21_00356. Due to ethical and legal restrictions, these data are not publicly available. Researchers can apply for access to Lifelines data through the Lifelines data access procedure (https://www.lifelines.nl/researcher/how-to-apply).

## Code availability

The source code implementing the PUMA framework and the analyses presented in this study is available at: https://github.com/GhorbaniF/PUMA-phenotypic-aging.

## Acknowledgements

The authors thank the participants of the Lifelines Cohort Study and the Lifelines research team for collecting, maintaining, and providing access to this valuable resource.

