## Supplementary for "PUMA: A Phenotypic Unsupervised Model of Aging Reveals Distinct Aging Dimensions"

### Supplementary Tables

**Supplementary Table 1.** Coding scheme used to identify incident disease cases using self-reported data available at both the second and third follow-up assessments.

| **Baseline assessment variable:**<br><br>**Are you currently diagnosed with cancer, diabetes, COPD, heart failure, stroke and Parkinson's disease (yes=1, No=2)** | **Second assessment variable:**<br><br>**Did you develop cancer, diabetes, COPD, heart failure, stroke and Parkinson's disease since the last time you filled in the lifeline questionnaire (yes=1, No=2)** | **Third assessment variable:**<br><br>**Did you develop cancer, diabetes, COPD, heart failure, stroke and Parkinson's disease since the last time you filled in the lifeline questionnaire (yes=1, No=2)** | **Incident disease case**<br><br>**(No=0, Yes=1)** |
|---|---|---|---|
| 1 | 1 | NA | 0 |
| 1 | 2 | NA | 0 |
| 1 | NA | 2 | 0 |
| 1 | NA | 1 | 0 |
| 1 | 2 | 2 | 0 |
| 1 | 1 | 1 | 0 |
| 1 | 2 | 1 | 0 |
| 1 | 1 | 2 | 0 |
| 1 | NA | NA | 0 |
| 2 | 2 | NA | 0 |
| 2 | 1 | NA | 1 |
| 2 | NA | 1 | 1 |
| 2 | NA | NA | NA |
| 2 | NA | 2 | 0 |
| 2 | 2 | 2 | 0 |
| 2 | 2 | 1 | 1 |
| 2 | 1 | 1 | 1 |
| 2 | 1 | 2 | 1 |

| | | | |
|---|---|---|---|
| NA | 2 | 2 | 0 |
| NA | NA | 2 | 0 |
| NA | 1 | 2 | 1 |
| NA | 2 | 1 | 1 |
| NA | 1 | 1 | 1 |
| NA | 1 | NA | 1 |
| NA | 2 | NA | 0 |
| NA | NA | 1 | 1 |
| NA | NA | NA | NA |

**Supplementary Table 2.** Coding scheme used to identify incident disease cases using self-reported data available at the second or third follow-up assessments.

| **Baseline assessment variable:** **Are you currently diagnosed with cancer, diabetes, COPD, heart failure, stroke and Parkinson's disease (yes=1, No=2)** | **Second or third assessment variable:** **Did you develop cancer, diabetes, COPD, heart failure, stroke and Parkinson's disease since the last time you filled in the lifeline questionnaire (yes=1, No=2)** | Incident disease case **(No=0, Yes=1)** |
|---|---|---|
| 1 | 1 | 0 |
| 1 | 2 | 0 |
| 1 | NA | 0 |
| 2 | 1 | 1 |
| 2 | 2 | 0 |
| 2 | NA | NA |
| NA | 1 | 1 |
| NA | 2 | 0 |
| NA | NA | NA |

**Supplementary Table 3. Definitions of Lifelines' subsections**

| Sub-section | Definition |
|---|---|
| Pets & Livestock | Exposure to pets and livestock |
| Exposure to smoking | Adult participants and parents of underage participants were asked whether they / their children were ever exposed to secondhand tobacco smoke |
| Informal care | Participants were asked whether they gave unpaid care to someone close (family/friend) because of long term limitations and/or health problems, excluding unpaid care of strangers (i.e. volunteer work) or the care for healthy children |
| Home environment (general) | Participants were asked several questions regarding the environment at home etc, heating, house moving, type of house, |
| Smoking and tobacco use | Participants were asked whether they (had) smoked any tobacco |
| Smoking derivatives | Adult Lifelines participants were asked whether they (had) smoked any tobacco. |
| Quality of Life (RAND) | The RAND-36 measures 'Health Related Quality of Life', i.e. how health impacts an individual's ability to function and his or her perceived well-being in physical, mental and social domains of life |
| Work absence | Absence from work was assessed in adult Lifelines participants |
| Treatment (general) | Lifelines participants were asked about the medical treatments they underwent in their lifetime, in addition to hospitalization, surgery, and medication |
| Household composition | Adult Lifelines participants were asked the various regarding their household composition etc, partner, divorce, who you are living with, number of people at home |
| Personal data | Questions from the person, etc, left or right handed, |
| Nationality & Ethnicity | Lifelines participants were asked the following questions about their experiences with cancer |
| Cancer | Experience with cancer |
| Cardiovascular diseases | Adult and elderly Lifelines participants were asked whether they suffered from various cardiovascular diseases. |
| Cardiovascular symptoms | Lifelines participants were asked whether they suffered from the undiagnosed symptoms associated with cardiovascular diseases. Etc, chest pain, edema |
| Diseases (other) | Lifelines participants were asked whether they suffered from any diseases other than the ones that were specified in the questionnaire(s). do or did you suffer from any other disorder that you have not mentioned yet? |
| Ear conditions | Lifelines participants were asked the following questions about health issues with their ears |
| Eye conditions (general) | Lifelines participants were asked the following general questions about their eyes and eye sight |
| Kidney & bladder diseases | Lifelines participants were asked whether they have had and/or were treated for a number of kidney & bladder disease. |

| | |
|---|---|
| Musculoskeletal conditions | Lifelines participants were asked whether they had the following muscle, joint, and/or bone diseases |
| Musculoskeletal symptoms | Lifelines participants were asked whether they suffered from the following undiagnosed symptoms (i.e. pain) in their musculoskeletal system. |
| Oral cavity conditions | Lifelines participants were asked the following questions about their oral health. have you been diagnosed with periodontitis by your/a dentist? |
| Respiratory diseases | Lifelines participants were asked whether they had and/or were treated for various respiratory diseases etc asthma and copd |
| Respiratory Problems (general) | Lifelines participants were asked whether they had any (undiagnosed) respiratory symptoms etc wheezing, shortensess of breath, coughing |
| Surgery | Lifelines participants were asked about any surgeries they have had |
| Medication | Lifelines participants were asked several questions about their medication use. |
| Stress (LDI) | The Long-term Difficulties Inventory (LDI) is a self-report questionnaire to measure long-term difficulties. It consists of 12 items referring to aspects of life, including housing, work, social relationships, free time, finances, health, school/study, and religion. Respondents have to indicate for each aspect how they experienced these aspects with respect to difficulty and stress in the last 12 months, on a three-point scale. |
| Personality (NEO) | The NEO-PI-R is a self-report instrument that measures the five most important domains of personality: neuroticism, extraversion, openness, agreeableness and conscientiousness |
| Stress (LTE) | The List of Threatening Experiences (LTE) is an instrument developed by Traolach Brugha and measures the occurrence of 12 major categories of stressful life events via self-report. The 12 major categories of stressful life events were selected for their established long-term consequences. The LTE is a 12-item self report questionnaire that measures stressful life events. For each response category, participants indicated whether or not they experienced each of these events in the last 12 months (yes=1/no=0). |
| Anthropometry (measured) | Height, weight, waist and hip circumference were assessed. |
| Anthropometry (self-reported) | Lifelines participants were asked to self-report information about their length and weight |
| ECG | Resting ECGs (section: physical state) were taken in Lifelines participants aged 13 years and older during: |
| Biosamples - blood | The following variables were assessed in the plasma of fresh blood samples collected from Lifelines participants. Blood parameters |
| Biosamples - urine | The following variables were assessed in fresh urine samples collected from Lifelines participants during etc, creatinine, albumin, |
| Employment status | The following questions were asked regarding the current employment status of adults |
| Contraception | Lifelines participants were asked about their use of contraceptives |

| | |
|---|---|
| Gynecological surgery | Adult female Lifelines participants were asked about any gynecological surgeries they had |
| Pregnancy outcome | Female Lifelines participants were asked whether they were or had ever been pregnant, and what the outcome of these pregnancies was. Etc, miscarriage, pregnancy, abortion, pregannacy duration |
| Birth (of participant) | Adult Lifelines participants were asked about their own birth (weight, hight, birth defect, how you were born, breast feed) |
| Menstruation | Adult and young female Lifelines participants were asked about menstruation and menstrual pain etc, start, duration, stop |
| Physical activity (SQUASH scores) | The Short questionnaire to Assess Health-enhancing physical activity (SQUASH) is a Dutch questionnaire developed by the Dutch National Institute of Public Health and the Environment to give an indication of habitual physical activity levels. |
| Personality sum scores | Calculated scores for several facets and five factor inventory (FFI) domains of the Personality (NEO) from participants aged 18-65. |
| Macronutrients (FFQ-h) [2] | Dietary intake of Lifelines participants aged 13 years and older was assessed with a food frequency questionnaire (FFQ) |
| Kidney/bladder diseases | Lifelines participants were asked whether they have had and/or were treated for a number of kidney & bladder diseases |
| Heart rate variability | Performed a validation of the ECG. |
| Family composition | The following questions were asked regarding family composition, i.e. the birth/existence of parents, siblings, children and partners of Lifelines participants. |

**Supplementary Table 4**. Key variables contributing to PC1-PC7, their partial correlation values, description based on the Lifelines' catalog and the Lifelines' subsection in which they are classified.

| PC | Number | Key variables | Partial correlation values | Description | Subsection |
|---|---|---|---|---|---|
| 1 | 1 | bird_childhood_adu_q_1_a | 0.332774 | was there a bird in your home during your first year of life? | Pets & livestock |
| | 2 | bird_childhood_adu_q_1_b | 0.302426 | was there a bird in your home when you were 1-4 years old? | Pets & livestock |
| | 3 | bird_childhood_adu_q_1_c | 0.288815 | was there a bird in your home when you were 5-15 years old? | Pets & livestock |
| | 4 | bird_presence_adu_q_1 | 0.441132 | do you have a bird? | Pets & livestock |
| | 5 | birth_defects_adu_q_1 | 0.313094 | did you have one or more birth defects at birth? | Birth (of participant) |

| | | | | | |
|---|---|---|---|---|---|
| | 6 | bodyweight_diet_adu_q_1 | 0.337465 | are you currently on a diet to lose weight? | Anthropometry (self-reported) |
| | 7 | bodyweight_shortdiet_adu_q_1 | 0.24186 | do you sometimes follow a diet for a short period of time (at least 1 day), during which you deliberately eat less than usual (ie slimming)? | Anthropometry (self-reported) |
| | 8 | dog_presence_adu_q_1 | 0.250109 | do you have a dog? | Pets & livestock |
| | 9 | ffqh_diet_adu_q_01 | 0.298432 | were you on a diet in the past month? | Macronutrients (FFQ-h) |
| | 10 | ffqh_diet_adu_q_03 | 0.241968 | were you on a diet based on certain beliefs/convictions (eg vegetarian or macrobiotic)? | Macronutrients (FFQ-h) |
| | 11 | guineapig_childhood_adu_q_1_a | 0.460333 | was there a guinea pig in your home during your first year of life? | Pets & livestock |
| | 12 | guineapig_childhood_adu_q_1_b | 0.406854 | was there a guinea pig in your home when you were 1-4 years old? | Pets & livestock |
| | 13 | guineapig_childhood_adu_q_1_c | 0.377634 | was there a guinea pig in your home when you were 5-15 years old? | Pets & livestock |
| | 14 | guineapig_presence_adu_q_1 | 0.436636 | do you have a guinea pig? | Pets & livestock |
| | 15 | ldi_religion_adu_q_12 | 0.263791 | faith, church or religion (e.g. doubt, conflicts with your minister) / to what extent did you experience difficulties and stress related to this aspect of your life in the past year? | Stress (LDI) |
| | 16 | ldi_school_adu_q_11 | 0.201951 | school/study (e.g. too difficult, cannot be combined with other tasks) / to what extent did you experience difficulties and stress related to this aspect of your life in the past year? | Stress (LDI) |
| | 17 | livingroom_heating_adu_q_1_a | 0.407366 | central heating / how is your living room heated? | Home environment (general) |
| | 18 | lte_pastyear_adu_q_01 | 0.34652 | you were severely ill, severely injured or a victim of violence / could you indicate whether you experienced this unpleasant event in the past year? | Stress (LTE) |
| | 19 | lte_pastyear_adu_q_03 | 0.265403 | a parent, child, brother, sister or partner died / could you indicate whether you experienced this unpleasant event in the past year? | Stress (LTE) |
| | 20 | lte_pastyear_adu_q_05 | 0.258526 | you and your partner split up / could you indicate whether you | Stress (LTE) |

| | | | | | |
|---|---|---|---|---|---|
| | | | | experienced this unpleasant event in the past year? | |
| | 21 | lte_pastyear_adu_q_06 | 0.287516 | you ended a long-term relationship with a good friend or relative / could you indicate whether you experienced this unpleasant event in the past year? | Stress (LTE) |
| | 22 | lte_pastyear_adu_q_07 | 0.313752 | you got into a serious problem with a good friend, relative or neighbour / could you indicate whether you experienced this unpleasant event in the past year? | Stress (LTE) |
| | 23 | lte_pastyear_adu_q_08 | 0.276626 | you lost your job and haven't been able to find work again / could you indicate whether you experienced this unpleasant event in the past year? | Stress (LTE) |
| | 24 | lte_pastyear_adu_q_09 | 0.299314 | you were fired / could you indicate whether you experienced this unpleasant event in the past year? | Stress (LTE) |
| | 25 | lte_pastyear_adu_q_10 | 0.349794 | you faced severe financial difficulties / could you indicate whether you experienced this unpleasant event in the past year? | Stress (LTE) |
| | 26 | lte_pastyear_adu_q_11 | 0.405036 | you got into trouble with the police or the law / could you indicate whether you experienced this unpleasant event in the past year? | Stress (LTE) |
| | 27 | lte_pastyear_adu_q_12 | 0.398238 | you lost money or valuables through theft or loss / could you indicate whether you experienced this unpleasant event in the past year? | Stress (LTE) |
| | 28 | lte_pastyear_adu_q_13 | 0.355854 | did any other important and serious negative events happen to you in the past year? | Stress (LTE) |
| | 29 | neonumber_competence_adu_c_1 | 0.302858 | number of valid items within competence facet | Personality sum scores |
| | 30 | neonumber_deliberation_adu_c_1 | 0.304104 | number of valid items within deliberation facet | Personality sum scores |
| | 31 | neonumber_excitement_adu_c_1 | 0.305276 | number of valid items within excitement-seeking facet | Personality sum scores |
| | 32 | neonumber_hostility_adu_c_1 | 0.307325 | number of valid items within anger-hostility facet | Personality sum scores |
| | 33 | neonumber_impulsivity_adu_c_1 | 0.304197 | number of valid items within impulsivity facet | Personality sum scores |

| | | | | | |
|---|---|---|---|---|---|
| | 34 | neonumber_neuroticism_adu_c_1 | 0.216355 | number of valid items within neuroticism domain | Personality sum scores |
| | 35 | neonumber_selfconsciousness_adu_c_1 | 0.306841 | number of valid items within self-consciousness facet | Personality sum scores |
| | 36 | neonumber_selfdiscipline_adu_c_1 | 0.307649 | number of valid items within self-discipline facet | Personality sum scores |
| | 37 | neonumber_vulnerability_adu_c_1 | 0.295653 | number of valid items within vulnerability facet | Personality sum scores |
| | 38 | smoke_father_adu_q_1 | 0.366822 | has your father ever smoked regularly during your childhood? | Exposure to smoke |
| | 39 | smoke_mother_adu_q_1 | 0.342964 | has your mother ever smoked regularly during your childhood? | Exposure to smoke |
| | 40 | squashsum_missing_adu_c_1 | 0.399273 | complete questionnaire missing | Physical activity (SQUASH scores) [2] |
| 2 | 1 | bird_presence_adu_q_1 | 0.21333 | do you have a bird? | Pets & livestock |
| | 2 | care_timespending_adu_q_1_a | 0.213895 | not applicable, i do not provide care for extended family members, friends or neighbors / on average how many hours per week do you spend caring for one or more extended family members, friends, or neighbors? | Informal care |
| | 3 | guineapig_presence_adu_q_1 | 0.200007 | do you have a guinea pig? | Pets & livestock |
| | 4 | inhouse_stepchildren_adu_q_1_v2 | 0.267252 | children of my partner that are not my own (stepchildren) / who are living in your house? (more than half the time) | Household composition |
| | 5 | ldi_freetime_adu_q_08 | 0.229132 | free time (e.g. too little or too much free time) / to what extent did you experience difficulties and stress related to this aspect of your life in the past year? | Stress (LDI) |
| | 6 | ldi_friends_adu_q_03 | 0.240859 | relationship with friends or acquaintances (e.g. quarrels, lack of support) / to what extent did you experience difficulties and stress related to this aspect of your life in the past year? | Stress (LDI) |
| | 7 | ldi_parents_adu_q_06 | 0.204503 | relationship with your parents (e.g. frequent conflicts, lack of acceptance) / to what extent did you experience difficulties and stress related to this aspect of your life in the past year? | Stress (LDI) |

| | | | | | |
|---|---|---|---|---|---|
| | 8 | ldi_relatives_adu_q_07 | 0.20701 | relationship with other relatives (e.g. conflicts, lack of acceptance) / to what extent did you experience difficulties and stress related to this aspect of your life in the past year? | Stress (LDI) |
| | 9 | living_assisted_adu_q_1 | 0.291963 | i live in an assisted living facility / what is your living situation? | Home environment (general) |
| | 10 | living_independent_adu_q_1 | 0.237713 | i live independently / what is your living situation? | Home environment (general) |
| | 11 | living_nursinghome_adu_q_1 | 0.316174 | i live in a nursing home / what is your living situation? | Home environment (general) |
| | 12 | living_residentialcare_adu_q_1 | 0.316227 | i live in a residential care home / what is your living situation? | Home environment (general) |
| | 13 | living_serviceflat_adu_q_1 | 0.291022 | i live in a service flat/sheltered housing complex / what is your living situation? | Home environment (general) |
| | 14 | living_withhelp_adu_q_1 | 0.28916 | i live independently with help / what is your living situation? | Home environment (general) |
| | 15 | lte_pastyear_adu_q_12 | 0.201406 | you lost money or valuables through theft or loss / could you indicate whether you experienced this unpleasant event in the past year? | Stress (LTE) |
| | 16 | neo_achievement_adu_q_82 | 0.288774 | i work hard to reach my goals | Personality (NEO) |
| | 17 | neo_achievement_adu_q_95 | 0.334875 | i strive to excel in everything i do | Personality (NEO) |
| | 18 | neo_activity_adu_q_81 | 0.218653 | i often feel like i'm bursting with energy | Personality (NEO) |
| | 19 | neo_activity_adu_q_94 | 0.3668 | i have a hectic life | Personality (NEO) |
| | 20 | neo_activity_adu_q_98 | 0.260929 | i am a very active person | Personality (NEO) |
| | 21 | neo_anxiety_adu_q_74 | 0.255099 | i rarely feel anxious or concerned | Personality (NEO) |
| | 22 | neo_anxiety_adu_q_79 | 0.219309 | i often feel tense and nervous | Personality (NEO) |

| | | | | | |
|---|---|---|---|---|---|
| | 23 | neo_anxiety_adu_q_92 | 0.292878 | i have fewer fears than most people | Personality (NEO) |
| | 24 | neo_depression_adu_q_66 | 0.263784 | i rarely feel lonely or sad | Personality (NEO) |
| | 25 | neo_depression_adu_q_77 | 0.257684 | i am rarely sad or depressed | Personality (NEO) |
| | 26 | neo_depression_adu_q_93 | 0.238855 | sometimes everything seems rather bleak and hopeless to me | Personality (NEO) |
| | 27 | neo_dutifulness_adu_q_72 | 0.470908 | sometimes i am not as reliable as i should be | Personality (NEO) |
| | 28 | neo_dutifulness_adu_q_86 | 0.218256 | when i make a promise, people can count on it that i will keep it | Personality (NEO) |
| | 29 | neo_excitement_adu_q_37 | 0.210491 | i like to be there where something is going on | Personality (NEO) |
| | 30 | neo_gregariousness_adu_q_75 | 0.244682 | i usually prefer to do things on my own | Personality (NEO) |
| | 31 | neo_order_adu_q_70 | 0.389624 | i keep my belongings neat and clean | Personality (NEO) |
| | 32 | neo_order_adu_q_76 | 0.330789 | i am not very systematic | Personality (NEO) |
| | 33 | neo_order_adu_q_84 | 0.433657 | i just cannot seem to get things sorted out | Personality (NEO) |
| | 34 | neo_positive_adu_q_78 | 0.367069 | i am not a cheerful optimist | Personality (NEO) |
| | 35 | neo_positive_adu_q_87 | 0.376228 | i don't really see myself as a happy and cheerful person | Personality (NEO) |
| | 36 | neo_positive_adu_q_91 | 0.29891 | i am a cheerful and lively person | Personality (NEO) |
| | 37 | neo_positive_adu_q_99 | 0.274775 | i laugh easily | Personality (NEO) |
| | 38 | neovalid_conscientiousness_adu_c_1 | 0.221284 | sum of all valid items within conscientiousness domain | Personality sum scores |
| | 39 | partner_deceased_adu_q_1 | 0.232548 | did you ever have a partner who passed away? | Household composition |
| | 40 | permanent_address_adu_q_1_a | 0.258221 | i've always had a permanent address / for how many years have you not had a permanent address? | Home environment (general) |
| | 41 | urems_blood_all_m_1 | 0.249158 | was the early morning spot urine sample obtained within 24h (before or after) the blood sample? | Biospecimen: Urine |

| | | | | | |
|---|---|---|---|---|---|
| | 42 | urems_ur24h_all_m_1 | 0.33471 | was the early morning spot urine sample obtained within 24h (before or after) the 24h urine sample? | Biospecimen: Urine |
| | 43 | uremsteststrip_blood_all_m_1 | 0.313939 | blood level in early morning spot urine sample according to test strip | Biospecimen: Urine |
| | 44 | uremsteststrip_glucose_all_m_1 | 0.278735 | glucose level in early morning spot urine sample according to test strip | Biospecimen: Urine |
| | 45 | uremsteststrip_ketone_all_m_1 | 0.29363 | ketone level in early morning spot urine sample according to test strip | Biospecimen: Urine |
| | 46 | uremsteststrip_leukocytes_all_m_1 | 0.290801 | leukocyte level in early morning spot urine sample according to test strip | Biospecimen: Urine |
| | 47 | uremsteststrip_nitrite_all_m_1 | 0.297711 | nitrite level in early morning spot urine sample according to test strip | Biospecimen: Urine |
| | 48 | uremsteststrip_ph_all_m_1 | 0.261263 | ph of early morning spot urine sample according to test strip | Biospecimen: Urine |
| | 49 | uremsteststrip_protein_all_m_1 | 0.298381 | protein level in early morning spot urine sample according to test strip | Biospecimen: Urine |
| 3 | 1 | absence_pastyear_adu_q_2_a | 0.225541 | in the past year, did you stay home from work because of an illness or problems for one or more periods of more than two weeks? (pregnancy is not considered an illness or problem) | Work absence |
| | 2 | employment_current_adu_q_1 | 0.201782 | do you do paid work, even if that is only for one or a few hours a week? | Employment status |
| | 3 | highcholesterol_presence_adu_q_1 | 0.20935 | have you ever been diagnosed with high cholesterol? | Cardiovascular diseases |
| | 4 | neo_anxiety_adu_q_88 | 0.239973 | i often worry about things that could go wrong | Personality (NEO) |
| | 5 | neo_dutifulness_adu_q_72 | 0.205402 | sometimes i am not as reliable as i should be | Personality (NEO) |
| | 6 | neo_gregariousness_adu_q_75 | 0.20568 | i usually prefer to do things on my own | Personality (NEO) |
| | 7 | neo_order_adu_q_84 | 0.205081 | i just cannot seem to get things sorted out | Personality (NEO) |
| | 8 | neonumber_competence_adu_c_1 | 0.324422 | number of valid items within competence facet | Personality sum scores |
| | 9 | neonumber_deliberation_adu_c_1 | 0.328992 | number of valid items within deliberation facet | Personality sum scores |

| | | | | | |
|---|---|---|---|---|---|
| | 10 | neonumber_excitement_adu_c_1 | 0.325364 | number of valid items within excitement-seeking facet | Personality sum scores |
| | 11 | neonumber_hostility_adu_c_1 | 0.331137 | number of valid items within anger-hostility facet | Personality sum scores |
| | 12 | neonumber_impulsivity_adu_c_1 | 0.326403 | number of valid items within impulsivity facet | Personality sum scores |
| | 13 | neonumber_neuroticism_adu_c_1 | 0.249383 | number of valid items within neuroticism domain | Personality sum scores |
| | 14 | neonumber_selfconsciousness_adu_c_1 | 0.327801 | number of valid items within self-consciousness facet | Personality sum scores |
| | 15 | neonumber_selfdiscipline_adu_c_1 | 0.332465 | number of valid items within self-discipline facet | Personality sum scores |
| | 16 | neonumber_vulnerability_adu_c_1 | 0.319243 | number of valid items within vulnerability facet | Personality sum scores |
| | 17 | rand_physical_adu_q_03_e | 0.206588 | climbing one flight of stairs / does your health now limit you in the following activities? | Quality of Life (RAND) |
| | 18 | rand_physical_adu_q_03_g | 0.208628 | walking more than a kilometer / does your health now limit you in the following activities? | Quality of Life (RAND) |
| | 19 | rand_physical_adu_q_03_h | 0.21696 | walking half a kilometer / does your health now limit you in the following activities? | Quality of Life (RAND) |
| | 20 | rand_physical_adu_q_03_i | 0.2146 | walking one hundred meters / does your health now limit you in the following activities? | Quality of Life (RAND) |
| | 21 | rmssd_ln_adu_c_1 | 0.202006 | log-transformed root mean square of successive differences (ms) | Heart rate variability [2] |
| | 22 | rmssd_lncorrected_adu_c_1 | 0.206618 | extreme value for log-transformed root mean square of successive differences corrected for heart rate (>5sd from mean) | Heart rate variability [2] |
| | 23 | sdnn_ln_adu_c_1 | 0.206411 | log-transformed standard deviation of interbeat interval (ms) | Heart rate variability [2] |
| | 24 | sdnn_lncorrected_adu_c_1 | 0.212857 | log-transformed standard deviation of interbeat interval corrected for heart rate (ms) | Heart rate variability [2] |
| | 25 | smoke_workspace_adu_q_1 | 0.209382 | do people smoke regularly in the space where you work? | Exposure to smoke |
| 4 | 1 | abortion_frequency_adu_q_1 | 0.213362 | how often have you had an abortion? | Pregnancy outcomes |

| | | | | | |
|---|---|---|---|---|---|
| | 2 | bodyweight_at18_adu_q_1 | 0.398784 | exactly (xxx kg) / for women: how much did you weigh when you were 18 years old? | Anthropometry (self-reported) |
| | 3 | bodyweight_at20_adu_q_1 | 0.349016 | for men: how much did you weigh when you were 20 years old? | Anthropometry (self-reported) |
| | 4 | carotid_stenosis_adu_q_1 | 0.23489 | have you ever been diagnosed with a narrowing in one or both carotid arteries? | Cardiovascular diseases |
| | 5 | cycle_current_adu_q_1 | 0.229571 | are you still menstruating? | Menstruation |
| | 6 | gp_pastyear_adu_q_1 | 0.236796 | gp / please fill in which of the health providers listed below you have contacted for yourself in the past 12 months | Treatment (general) |
| | 7 | hdlchol_result_all_m_1 | 0.22998 | hdl cholesterol in lithium heparin tube (mmol/l) | Biospecimen: Blood |
| | 8 | hormonal_contraception_adu_q_1 | 0.50656 | have you ever used hormonal contraception? | Contraception |
| | 9 | hormonal_contraception_adu_q_1_b | 0.301748 | did you use hormonal contraception in the past month? | Contraception |
| | 10 | hormonal_treatment_adu_q_1 | 0.583339 | have you ever received hormonal treatment for a reason other than contraception? | Medication |
| | 11 | hysterectomy_presence_adu_q_1 | 0.64543 | have you had a hysterectomy? | Gynecological surgery |
| | 12 | kidneydisease_diagnosis_adu_q_1 | 0.217423 | have you been diagnosed with a kidney disease? | Kidney / bladder diseases |
| | 13 | kidneysurgery_none_adu_q_1 | 0.214656 | no / have you ever had kidney surgery? | Surgery |
| | 14 | measurement_position_all_m_1 | 0.248046 | standing or sitting | Anthropometry (measured) |
| | 15 | menarche_age_adu_q_1 | 0.520543 | how old were you when you had your first menstruation? | Menstruation |
| | 16 | nodisorder_general_adu_q_1 | 0.214818 | none of the disorders / could you indicate which of the following disorders you have (had)? | Diseases (other) |
| | 17 | oophorectomy_double_adu_q_1 | 0.646879 | have you had two ovaries removed? | Gynecological surgery |
| | 18 | oophorectomy_single_adu_q_1 | 0.647261 | have you had one ovary removed? | Gynecological surgery |

| | | | | | |
|---|---|---|---|---|---|
| | 19 | partner_gender_adu_q_1 | 0.232902 | what is your partner's gender? | Household composition |
| | 20 | physiotherapist_pastyear_adu_q_1 | 0.272014 | physical therapist / please fill in which of the health providers listed below you have contacted for yourself in the past 12 months | Treatment (general) |
| | 21 | pregnancy_current_adu_q_1 | 0.429668 | are you currently pregnant? | Pregnancy outcomes |
| | 22 | rand_physical_adu_q_03_a | 0.298657 | vigorous activities, such as running, lifting heavy objects, participating in strenuous sports / does your health now limit you in the following activities? | Quality of Life (RAND) |
| | 23 | rand_physical_adu_q_03_b | 0.273184 | moderate activities, such as moving a table, pushing a vacuum cleaner, bicycling / does your health now limit you in the following activities? | Quality of Life (RAND) |
| | 24 | rand_physical_adu_q_03_c | 0.268167 | lifting or carrying groceries / does your health now limit you in the following activities? | Quality of Life (RAND) |
| | 25 | rand_physical_adu_q_03_d | 0.264043 | climbing several flights of stairs / does your health now limit you in the following activities? | Quality of Life (RAND) |
| | 26 | rand_physical_adu_q_03_e | 0.252366 | climbing one flight of stairs / does your health now limit you in the following activities? | Quality of Life (RAND) |
| | 27 | rand_physical_adu_q_03_f | 0.270529 | bending, kneeling or stooping / does your health now limit you in the following activities? | Quality of Life (RAND) |
| | 28 | rand_physical_adu_q_03_g | 0.256303 | walking more than a kilometer / does your health now limit you in the following activities? | Quality of Life (RAND) |
| | 29 | rand_physical_adu_q_03_h | 0.256772 | walking half a kilometer / does your health now limit you in the following activities? | Quality of Life (RAND) |
| | 30 | rand_physical_adu_q_03_i | 0.276722 | walking one hundred meters / does your health now limit you in the following activities? | Quality of Life (RAND) |
| | 31 | rand_physical_adu_q_03_j | 0.292311 | bathing or dressing yourself / does your health now limit you in the following activities? | Quality of Life (RAND) |
| | 32 | specialist_pastyear_adu_q_1 | 0.27275 | medical specialist / please fill in which of the health providers listed below you have contacted for yourself in the past 12 months | Treatment (general) |

| | | | | | |
|---|---|---|---|---|---|
| | 33 | urinetest_cystitis_adu_q_1 | 0.403078 | (signs of) cystitis or kidney stones / do you know the outcome of these urine tests? | Kidney / bladder diseases |
| 5 | 1 | amputation_leg_adu_q_1 | 0.369651 | was a (part of a) foot/leg amputated? | Musculoskeletal conditions |
| | 2 | aneurysm_diagnosis_adu_q_1 | 0.372957 | were you ever diagnosed with a dilatation of the aorta (aortic aneurysm)? | Cardiovascular diseases |
| | 3 | angioplasty_bypass_adu_q_1 | 0.320669 | have you ever had a balloon angioplasty (stretching of artery with balloon) and/or bypass surgery? | Surgery |
| | 4 | bird_presence_adu_q_1 | 0.211939 | do you have a bird? | Pets & livestock |
| | 5 | bodyweight_at18_adu_q_1 | 0.289299 | for women: how much did you weigh when you were 18 years old? | Anthropometry (self-reported) |
| | 6 | cancer_lifetime_adu_q_1 | 0.341499 | do you have cancer or have you had cancer? | Cancer |
| | 7 | carotid_stenosis_adu_q_1 | 0.367174 | have you ever been diagnosed with a narrowing in one or both carotid arteries? | Cardiovascular diseases |
| | 8 | copd_presence_adu_q_1 | 0.295647 | do you have copd, emphysema or chronic bronchitis? | Respiratory diseases |
| | 9 | coughing_presence_adu_q_1 | 0.279033 | have you ever woken up from a coughing fit? | Respiratory problems (general) |
| | 10 | coughing_winter_adu_q_1 | 0.36766 | do you usually cough in the wintertime when getting up? | Respiratory problems (general) |
| | 11 | coughing_winter_adu_q_2 | 0.323425 | do you usually cough in wintertime? | Respiratory problems (general) |
| | 12 | disturbedkidney_diagnosis_adu_q_1 | 0.439164 | have you ever been diagnosed with a disturbed kidney function? | Kidney / bladder diseases |
| | 13 | dyspnea_lying_adu_q_1 | 0.354738 | do you become short of breath when you lie (too) flat? | Respiratory problems (general) |
| | 14 | dyspnea_night_adu_q_1 | 0.297807 | do you at times wake up at night with shortness of breath? | Respiratory problems (general) |
| | 15 | dyspnea_presence_adu_q_1 | 0.205947 | have you ever had an attack of shortness of breath during daytime while at rest? | Respiratory problems (general) |

| | | | | | |
|---|---|---|---|---|---|
| | 16 | dyspnea_presence_adu_q_1_b | 0.281539 | have you ever woken up from an attack of shortness of breath? | Respiratory problems (general) |
| | 17 | edema_presence_adu_q_1 | 0.446322 | do you at times have swollen ankles (edema), or does fluid accumulate elsewhere? | Cardiovascular symptoms |
| | 18 | eyesight_limitations_adu_q_1 | 0.27189 | are you limited by problems with your eyesight in daily life? | Eye conditions (general) |
| | 19 | fasting_participant_all_q_1 | 0.206248 | has the participant been fasting? | Biospecimen: Blood |
| | 20 | functionloss_presence_adu_q_1 | 0.334136 | have you ever had a sudden loss of function which you later regained (such as loss of strength, sensory disturbance, blindness in one eye, slurred speech)? | Cardiovascular symptoms |
| | 21 | guineapig_childhood_adu_q_1_a | 0.231534 | was there a guinea pig in your home during your first year of life? | Pets & livestock |
| | 22 | guineapig_presence_adu_q_1 | 0.223054 | do you have a guinea pig? | Pets & livestock |
| | 23 | hearing_aid_adu_q_1 | 0.429256 | do you need a hearing aid? | Ear conditions |
| | 24 | hearing_limitations_adu_q_1 | 0.393916 | are you limited by problems with your hearing in daily life? | Ear conditions |
| | 25 | heartattack_presence_adu_q_1 | 0.311225 | have you ever had a heart attack? | Cardiovascular diseases |
| | 26 | hormonal_contraception_adu_q_1 | 0.294079 | have you ever used hormonal contraception? | Contraception |
| | 27 | hormonal_treatment_adu_q_1 | 0.337509 | have you ever received hormonal treatment for a reason other than contraception? | Medication |
| | 28 | hysterectomy_presence_adu_q_1 | 0.35977 | have you had a hysterectomy? | Gynecological surgery |
| | 29 | joints_pain_adu_q_1_a | 0.339618 | do you regularly (several times a week) have pain in the joints of your hands? | Musculoskeletal symptoms |
| | 30 | joints_pain_adu_q_1_b | 0.317963 | do you regularly (several times a week) have pain in the joints of your feet? | Musculoskeletal symptoms |
| | 31 | joints_stiffness_adu_q_1_a | 0.333743 | do you regularly (several times a week) have stiffness in the joints of your hands? | Musculoskeletal symptoms |

| | | | | | |
|---|---|---|---|---|---|
| | 32 | joints_stiffness_adu_q_1_b | 0.309081 | do you regularly (several times a week) have stiffness in the joints of your feet? | Musculoskeletal symptoms |
| | 33 | kidney_disorders_adu_q_1_a | 0.444122 | no kidney disorder / have you had one or more of the following kidney disorders? | Kidney / bladder diseases |
| | 34 | kidneydisease_diagnosis_adu_q_1 | 0.260258 | have you been diagnosed with a kidney disease? | Kidney / bladder diseases |
| | 35 | kidneyproblem_bloodtest_adu_q_1 | 0.2921 | was your blood ever tested for a possible kidney problem? | Kidney / bladder diseases |
| | 36 | kidneysurgery_none_adu_q_1 | 0.421387 | no / have you ever had kidney surgery? | Surgery |
| | 37 | lte_pastyear_adu_q_01 | 0.204515 | you were severely ill, severely injured or a victim of violence / could you indicate whether you experienced this unpleasant event in the past year? | Stress (LTE) |
| | 38 | lte_pastyear_adu_q_09 | 0.233525 | you were fired / could you indicate whether you experienced this unpleasant event in the past year? | Stress (LTE) |
| | 39 | lte_pastyear_adu_q_11 | 0.231569 | you got into trouble with the police or the law / could you indicate whether you experienced this unpleasant event in the past year? | Stress (LTE) |
| | 40 | lte_pastyear_adu_q_12 | 0.201616 | you lost money or valuables through theft or loss / could you indicate whether you experienced this unpleasant event in the past year? | Stress (LTE) |
| | 41 | menarche_age_adu_q_1 | 0.214003 | how old were you when you had your first menstruation? | Menstruation |
| | 42 | oophorectomy_double_adu_q_1 | 0.366352 | have you had two ovaries removed? | Gynecological surgery |
| | 43 | oophorectomy_single_adu_q_1 | 0.363742 | have you had one ovaries removed? | Gynecological surgery |
| | 44 | phlegm_winter_adu_q_1 | 0.360153 | do you usually cough up phlegm immediately after getting up in wintertime? | Respiratory problems (general) |
| | 45 | phlegm_winter_adu_q_2 | 0.355008 | do you usually cough up phlegm during daytime or at night in wintertime? | Respiratory problems (general) |

| | | | | | |
|---|---|---|---|---|---|
| | 46 | pregnancy_current_adu_q_1 | 0.217684 | are you currently pregnant? | Pregnancy outcomes |
| | 47 | rand_emotional_adu_q_09_f | 0.208786 | have you felt downhearted and blue? / how much of the time during the past 4 weeks | Quality of Life (RAND) |
| | 48 | rand_limitations_adu_q_04_a | 0.216323 | cut down the amount of time you spent on work or other activities / during the past 4 weeks, have you had any of the following problems with your work or other regular daily activities as a result of your physical health? | Quality of Life (RAND) |
| | 49 | rand_limitations_adu_q_04_b | 0.218806 | accomplished less than you would like / during the past 4 weeks, have you had any of the following problems with your work or other regular daily activities as a result of your physical health? | Quality of Life (RAND) |
| | 50 | rand_limitations_adu_q_04_c | 0.207867 | were limited in the kind of work or other activities / during the past 4 weeks, have you had any of the following problems with your work or other regular daily activities as a result of your physical health? | Quality of Life (RAND) |
| | 51 | rand_limitations_adu_q_04_d | 0.217442 | had difficulty performing the work or other activities (for example, it took extra effort) / during the past 4 weeks, have you had any of the following problems with your work or other regular daily activities as a result of your physical health? | Quality of Life (RAND) |
| | 52 | smoking_current_adu_q_1 | 0.231424 | do you smoke now, or have you smoked in the past month? | Smoking & tobacco use |
| | 53 | smoking_current_adu_qc_1 | 0.228195 | do you smoke now, or have you smoked in the past month? | Smoking derivatives [2] |
| | 54 | squashsum_missing_adu_c_1 | 0.220259 | complete questionnaire missing | Physical activity (SQUASH scores) [2] |
| | 55 | stroke_presence_adu_q_1 | 0.334839 | have you ever had a stroke? | Cardiovascular diseases |
| | 56 | thyroid_medication_adu_q_1_a | 0.359396 | do you currently use medication for an overactive or underactive thyroid? | Medication |
| | 57 | thyroid_medication_adu_q_1_b | 0.404758 | have you used medication for an overactive or underactive thyroid in the past? | Medication |

| | | | | | |
|---|---|---|---|---|---|
| | 58 | ur24h_missed_all_m_1 | 0.255124 | are any urinations for the 24h urine sample missed? | Biospecimen: Urine |
| | 59 | urinetest_cystitis_adu_q_1 | 0.205423 | (signs of) cystitis or kidney stones / do you know the outcome of these urine tests? | Kidney / bladder diseases |
| | 60 | woundhealing_feet_adu_q_1 | 0.422328 | have you ever had poorly healing wounds on your feet? | Musculoskeletal symptoms |
| 6 | 1 | basoconc_result_all_m_1 | 0.245182 | basophilic granulocytes in k2-edta tube (10e9/l) | Biospecimen: Blood |
| | 2 | bird_childhood_adu_q_1_a | 0.200842 | was there a bird in your home during your first year of life? | Pets & livestock |
| | 3 | birthplace_country_adu_q_1 | 0.225561 | in what country were you born? | Nationality & ethnicity |
| | 4 | birthplace_father_fam_q_1 | 0.227835 | in what country was your (biological) father born? | Nationality & ethnicity |
| | 5 | birthplace_mother_fam_q_1 | 0.23464 | in what country was your (biological) mother born? | Nationality & ethnicity |
| | 6 | bodyweight_at18_adu_q_1 | 0.225656 | for women: how much did you weigh when you were 18 years old? | Anthropometry (self-reported) |
| | 7 | bodyweight_at20_adu_q_1 | 0.211891 | for men: how much did you weigh when you were 20 years old? | Anthropometry (self-reported) |
| | 8 | childbirth_currentpartner_fam_q_1_02 | 0.235205 | gender child (1) / biological children with your current partner | Family composition |
| | 9 | cigarillos_frequency_adu_c_2 | 0.286547 | current number of cigarillos per day | Smoking derivatives [2] |
| | 10 | cigars_frequency_adu_c_2 | 0.546004 | current number of cigars per day | Smoking derivatives [2] |
| | 11 | current_smoker_adu_c_2 | 0.521753 | current smoker | Smoking derivatives [2] |
| | 12 | ecg_reviewed_all_e_1 | 0.369121 | is the ecg manually reviewed by a doctor? | ECG |

| | | | | | |
|---|---|---|---|---|---|
| | 13 | ecg_toreview_all_e_1 | 0.370872 | is ecg to be manually reviewed by a doctor? | ECG |
| | 14 | ever_smoker_adu_c_2 | 0.55161 | ever smoker | Smoking derivatives [2] |
| | 15 | ex_smoker_adu_c_2 | 0.503547 | ex smoker | Smoking derivatives [2] |
| | 16 | guineapig_childhood_adu_q_1_b | 0.207422 | was there a guinea pig in your home when you were 1-4 years old? | Pets & livestock |
| | 17 | handedness_leftright_adu_q_1 | 0.219918 | are you left or right handed? | Personal data |
| | 18 | hdlchol_result_all_m_1 | 0.239608 | hdl cholesterol in lithium heparin tube (mmol/l) | Biospecimen: Blood |
| | 19 | hormonal_contraception_adu_q_1 | 0.329787 | have you ever used hormonal contraception? | Contraception |
| | 20 | hormonal_treatment_adu_q_1 | 0.357748 | have you ever received hormonal treatment for a reason other than contraception? | Medication |
| | 21 | hysterectomy_presence_adu_q_1 | 0.361206 | have you had a hysterectomy? | Gynecological surgery |
| | 22 | kidneydisease_diagnosis_adu_q_1 | 0.216771 | have you been diagnosed with a kidney disease? | Kidney / bladder diseases |
| | 23 | menarche_age_adu_q_1 | 0.274752 | how old were you when you had your first menstruation? | Menstruation |
| | 24 | never_smoker_adu_c_1 | 0.518889 | never smoker | Smoking derivatives [2] |
| | 25 | oophorectomy_double_adu_q_1 | 0.365872 | have you had two ovaries removed? | Gynecological surgery |
| | 26 | oophorectomy_single_adu_q_1 | 0.367604 | have you had one ovaries removed? | Gynecological surgery |
| | 27 | partner_gender_adu_q_1 | 0.213088 | what is your partner's gender? | Household composition |

| | | | | | |
|---|---|---|---|---|---|
| | 28 | pipetobacco_frequency_adu_c_2 | 0.402847 | current grams of pipe tobacco per day | Smoking derivatives [2] |
| | 29 | pregnancy_current_adu_q_1 | 0.202873 | are you currently pregnant? | Pregnancy outcomes |
| | 30 | recent_starter_adu_c_2 | 0.420872 | recent starter | Smoking derivatives [2] |
| | 31 | rmssd_ln_adu_c_1 | 0.238426 | log-transformed root mean square of successive differences (ms) | Heart rate variability [2] |
| | 32 | rmssd_lncorrected_adu_c_1 | 0.241274 | extreme value for log-transformed root mean square of successive differences corrected for heart rate (>5sd from mean) | Heart rate variability [2] |
| | 33 | sdnn_ln_adu_c_1 | 0.243558 | log-transformed standard deviation of interbeat interval (ms) | Heart rate variability [2] |
| | 34 | sdnn_lncorrected_adu_c_1 | 0.247949 | log-transformed standard deviation of interbeat interval corrected for heart rate (ms) | Heart rate variability [2] |
| | 35 | smoking_current_adu_q_1 | 0.384505 | do you smoke now, or have you smoked in the past month? | Smoking & tobacco use |
| | 36 | smoking_current_adu_qc_1 | 0.369627 | do you smoke now, or have you smoked in the past month? | Smoking derivatives [2] |
| | 37 | smoking_duration_adu_c_2 | 0.234679 | duration of smoking in years | Smoking derivatives [2] |
| | 38 | smoking_fullyear_adu_q_1 | 0.532043 | have you ever smoked for a full year? | Smoking & tobacco use |
| | 39 | smoking_fullyear_adu_qc_1 | 0.476097 | have you ever smoked for a full year? | Smoking derivatives [2] |
| | 40 | smoking_habit_adu_c_2 | 0.538643 | smoking habits | Smoking derivatives [2] |
| | 41 | smoking_inhale_adu_q_1 | 0.586365 | do you inhale or have you inhaled while smoking? | Smoking & tobacco use |
| | 42 | smoking_stopped_adu_q_1 | 0.563171 | have you stopped smoking? | Smoking & tobacco use |
| | 43 | smoking_stopped_adu_qc_1 | 0.562537 | have you stopped smoking? | Smoking derivatives [2] |

| | | | | | |
|---|---|---|---|---|---|
| | 44 | smoking_type_adu_q_1_01 | 0.588844 | type of tobacco 1 / how much have you smoked up until now? | Smoking & tobacco use |
| | 45 | smoking_type_adu_qc_1_01 | 0.568183 | type of tabacco smoked during this period 1 / how much have you smoked up until now? | Smoking derivatives [2] |
| | 46 | ur24h_missed_all_m_1 | 0.249026 | are any urinations for the 24h urine sample missed? | Biospecimen: Urine |
| 7 | 1 | basoconc_result_all_m_1 | 0.240842 | basophilic granulocytes in k2-edta tube (10e9/l) | Biospecimen: Blood |
| | 2 | basoperc_result_all_m_1 | 0.314577 | basophilic granulocytes in k2-edta tube (%) | Biospecimen: Blood |
| | 3 | bird_childhood_adu_q_1_a | 0.293458 | was there a bird in your home during your first year of life? | Pets & livestock |
| | 4 | bird_childhood_adu_q_1_b | 0.282013 | was there a bird in your home when you were 1-4 years old? | Pets & livestock |
| | 5 | bird_childhood_adu_q_1_c | 0.259663 | was there a bird in your home when you were 5-15 years old? | Pets & livestock |
| | 6 | bird_presence_adu_q_1 | 0.301347 | do you have a bird? | Pets & livestock |
| | 7 | bodyweight_diet_adu_q_1 | 0.213679 | are you currently on a diet to lose weight? | Anthropometry (self-reported) |
| | 8 | childbirth_currentpartner_fam_q_1_02 | 0.233154 | gender child (1) / biological children with your current partner | Family composition |
| | 9 | cigarillos_frequency_adu_c_2 | 0.273711 | current number of cigarillos per day | Smoking derivatives [2] |
| | 10 | cigars_frequency_adu_c_2 | 0.531153 | current number of cigars per day | Smoking derivatives [2] |
| | 11 | current_smoker_adu_c_2 | 0.574726 | current smoker | Smoking derivatives [2] |
| | 12 | ever_smoker_adu_c_2 | 0.631735 | ever smoker | Smoking derivatives [2] |
| | 13 | ex_smoker_adu_c_2 | 0.620203 | ex smoker | Smoking derivatives [2] |
| | 14 | ffqh_diet_adu_q_01 | 0.209793 | were you on a diet in the past month? | Macronutrients (FFQ-h) |
| | 15 | guineapig_childhood_adu_q_1_a | 0.305875 | was there a guinea pig in your home during your first year of life? | Pets & livestock |
| | 16 | guineapig_childhood_adu_q_1_b | 0.318531 | was there a guinea pig in your home when you were 1-4 years old? | Pets & livestock |
| | 17 | guineapig_childhood_adu_q_1_c | 0.296133 | was there a guinea pig in your home when you were 5-15 years old? | Pets & livestock |
| | 18 | guineapig_presence_adu_q_1 | 0.280724 | do you have a guinea pig? | Pets & livestock |
| | 19 | hearing_limitations_adu_q_1 | 0.224203 | are you limited by problems with your hearing in daily life? | Ear conditions |
| | 20 | ldi_religion_adu_q_12 | 0.201703 | faith, church or religion (e.g. doubt, conflicts with your minister) / to what extent did you experience difficulties and stress related to this aspect of your life in the past year? | Stress (LDI) |

| | | | | | |
|---|---|---|---|---|---|
| | 21 | livingroom_heating_adu_q_1_a | 0.261836 | central heating / how is your living room heated? | Home environment (general) |
| | 22 | lte_pastyear_adu_q_09 | 0.242765 | you were fired / could you indicate whether you experienced this unpleasant event in the past year? | Stress (LTE) |
| | 23 | lte_pastyear_adu_q_12 | 0.208465 | you lost money or valuables through theft or loss / could you indicate whether you experienced this unpleasant event in the past year? | Stress (LTE) |
| | 24 | never_smoker_adu_c_1 | 0.560876 | never smoker | Smoking derivatives [2] |
| | 25 | periodontitis_diagnosis_adu_q_1 | 0.227299 | have you been diagnosed with periodontitis by your/a dentist? | Oral cavity conditions |
| | 26 | pipetobacco_frequency_adu_c_2 | 0.413114 | current grams of pipe tobacco per day | Smoking derivatives [2] |
| | 27 | rand_physical_adu_q_03_a | 0.221758 | vigorous activities, such as running, lifting heavy objects, participating in strenuous sports / does your health now limit you in the following activities? | Quality of Life (RAND) |
| | 28 | rand_physical_adu_q_03_b | 0.267317 | moderate activities, such as moving a table, pushing a vacuum cleaner, bicycling / does your health now limit you in the following activities? | Quality of Life (RAND) |
| | 29 | rand_physical_adu_q_03_c | 0.267837 | lifting or carrying groceries / does your health now limit you in the following activities? | Quality of Life (RAND) |
| | 30 | rand_physical_adu_q_03_d | 0.237028 | climbing several flights of stairs / does your health now limit you in the following activities? | Quality of Life (RAND) |
| | 31 | rand_physical_adu_q_03_e | 0.257795 | climbing one flight of stairs / does your health now limit you in the following activities? | Quality of Life (RAND) |
| | 32 | rand_physical_adu_q_03_f | 0.269992 | bending, kneeling or stooping / does your health now limit you in the following activities? | Quality of Life (RAND) |
| | 33 | rand_physical_adu_q_03_g | 0.232471 | walking more than a kilometer / does your health now limit you in the following activities? | Quality of Life (RAND) |
| | 34 | rand_physical_adu_q_03_h | 0.243263 | walking half a kilometer / does your health now limit you in the following activities? | Quality of Life (RAND) |
| | 35 | rand_physical_adu_q_03_i | 0.263088 | walking one hundred meters / does your health now limit you in the following activities? | Quality of Life (RAND) |
| | 36 | rand_physical_adu_q_03_j | 0.286511 | bathing or dressing yourself / does your health now limit you in the following activities? | Quality of Life (RAND) |
| | 37 | recent_starter_adu_c_2 | 0.408694 | recent starter | Smoking derivatives [2] |
| | 38 | smoke_workspace_adu_q_1 | 0.276879 | do people smoke regularly in the space where you work? | Exposure to smoke |
| | 39 | smoking_current_adu_q_1 | 0.356909 | do you smoke now, or have you smoked in the past month? | Smoking & tobacco use |

| | | | | | |
|---|---|---|---|---|---|
| | 40 | smoking_current_adu_qc_1 | 0.344562 | do you smoke now, or have you smoked in the past month? | Smoking derivatives [2] |
| | 41 | smoking_duration_adu_c_2 | 0.257853 | duration of smoking in years | Smoking derivatives [2] |
| | 42 | smoking_fullyear_adu_q_1 | 0.556311 | have you ever smoked for a full year? | Smoking & tobacco use |
| | 43 | smoking_fullyear_adu_qc_1 | 0.486694 | have you ever smoked for a full year? | Smoking derivatives [2] |
| | 44 | smoking_habit_adu_c_2 | 0.599504 | smoking habits | Smoking derivatives [2] |
| | 45 | smoking_inhale_adu_q_1 | 0.643574 | do you inhale or have you inhaled while smoking? | Smoking & tobacco use |
| | 46 | smoking_stopped_adu_q_1 | 0.627715 | have you stopped smoking? | Smoking & tobacco use |
| | 47 | smoking_stopped_adu_qc_1 | 0.625865 | have you stopped smoking? | Smoking derivatives [2] |
| | 48 | smoking_type_adu_q_1_01 | 0.65469 | type of tobacco 1 / how much have you smoked up until now? | Smoking & tobacco use |
| | 49 | smoking_type_adu_qc_1_01 | 0.627178 | type of tabacco smoked during this period 1 / how much have you smoked up until now? | Smoking derivatives [2] |
| | 50 | urinetest_cystitis_adu_q_1 | 0.208653 | (signs of) cystitis or kidney stones / do you know the outcome of these urine tests? | Kidney / bladder diseases |

**Supplementary Table 5**. Enrichment scores (ES), Normalized enrichment scores (NES), and their significance across all phenotypic aging dimensions and diseases.

| Disease | PC | High/Low scores | ES | NES | FDR-adjusted P-value |
|---|---|---|---|---|---|
| Cancer | PC1 | High | 0.299493 | 1.317958 | 0.001866 |
| | PC2 | High | 0.067507 | 0.503829 | 1 |
| | PC3 | High | 0.170756 | 1.204882 | 0.029242 |
| | PC4 | High | 0.071191 | 0.632946 | 1 |
| | PC5 | High | 0.248048 | 1.279376 | 0.001575 |
| | PC6 | High | 0.199211 | 1.099312 | 0.157184 |
| | PC7 | High | 0.20576 | 1.206174 | 0.006399 |
| | PC1 | Low | -0.17875 | -0.75479 | 1 |
| | PC2 | Low | -0.12799 | -1.78407 | 0.00056 |
| | PC3 | Low | -0.18226 | -0.98716 | 0.893266 |
| | PC4 | Low | -0.078 | -1.44792 | 0.009926 |
| | PC5 | Low | -0.14659 | -1.19362 | 0.098622 |
| | PC6 | Low | -0.1208 | -0.94825 | 1 |
| | PC7 | Low | -0.10715 | -0.86285 | 1 |
| Diabetes | PC1 | High | 0.466797 | 2.030161 | 0.00056 |
| | PC2 | High | 0.086971 | 0.646541 | 1 |
| | PC3 | High | 0.191227 | 1.341992 | 0.006299 |
| | PC4 | High | 0.121488 | 1.066586 | 0.518192 |

| | | | | | |
|---|---|---|---|---|---|
| | PC5 | High | 0.399952 | 2.025139 | 0.00056 |
| | PC6 | High | 0.314086 | 1.722072 | 0.00056 |
| | PC7 | High | 0.273343 | 1.590465 | 0.00056 |
| | PC1 | Low | -0.04648 | -0.1982 | 1 |
| | PC2 | Low | -0.1203 | -1.62191 | 0.001866 |
| | PC3 | Low | -0.22731 | -1.22173 | 0.017148 |
| | PC4 | Low | -0.14781 | -2.6712 | 0.00056 |
| | PC5 | Low | -0.08482 | -0.68991 | 1 |
| | PC6 | Low | -0.03845 | -0.29876 | 1 |
| | PC7 | Low | -0.07216 | -0.5767 | 1 |
| COPD | PC1 | High | 0.526081 | 1.805363 | 0.00056 |
| | PC2 | High | 0.127952 | 0.998598 | 0.804418 |
| | PC3 | High | 0.149244 | 1.016392 | 0.740806 |
| | PC4 | High | 0.180509 | 1.828172 | 0.00056 |
| | PC5 | High | 0.451661 | 1.718985 | 0.00056 |
| | PC6 | High | 0.284337 | 1.520422 | 0.00056 |
| | PC7 | High | 0.305095 | 1.430331 | 0.00056 |
| | PC1 | Low | -0.03476 | -0.22844 | 1 |
| | PC2 | Low | -0.0916 | -1.0281 | 0.736555 |
| | PC3 | Low | -0.28516 | -1.67014 | 0.00056 |
| | PC4 | Low | -0.06811 | -0.85073 | 1 |
| | PC5 | Low | -0.04487 | -0.5822 | 1 |
| | PC6 | Low | -0.04355 | -0.34667 | 1 |
| | PC7 | Low | -0.02707 | -0.3013 | 1 |
| Heart Failure | PC1 | High | 0.396734 | 1.247345 | 0.014607 |
| | PC2 | High | 0.127525 | 1.028173 | 0.731427 |
| | PC3 | High | 0.144 | 1.050224 | 0.634405 |
| | PC4 | High | 0.179993 | 1.482887 | 0.00221 |
| | PC5 | High | 0.331564 | 1.225219 | 0.022174 |
| | PC6 | High | 0.25074 | 1.21171 | 0.038396 |
| | PC7 | High | 0.170532 | 0.868554 | 1 |
| | PC1 | Low | -0.07113 | -0.62974 | 1 |
| | PC2 | Low | -0.06496 | -0.66813 | 1 |
| | PC3 | Low | -0.18368 | -1.08622 | 0.462954 |
| | PC4 | Low | -0.04754 | -0.71337 | 1 |
| | PC5 | Low | -0.10159 | -1.19679 | 0.384854 |
| | PC6 | Low | -0.05148 | -0.49242 | 1 |
| | PC7 | Low | -0.14639 | -1.37872 | 0.029242 |
| Stroke | PC1 | High | 0.465689 | 1.564786 | 0.00056 |
| | PC2 | High | 0.081962 | 0.641912 | 1 |
| | PC3 | High | 0.169229 | 1.135859 | 0.46586 |
| | PC4 | High | 0.170402 | 1.438889 | 0.061401 |
| | PC5 | High | 0.329715 | 1.256921 | 0.098622 |

| | | | | | |
|---|---|---|---|---|---|
| | PC6 | High | 0.238366 | 1.143595 | 0.320494 |
| | PC7 | High | 0.209611 | 1.007564 | 0.786614 |
| | PC1 | Low | -0.04451 | -0.32694 | 1 |
| | PC2 | Low | -0.13048 | -1.2765 | 0.289204 |
| | PC3 | Low | -0.18392 | -1.11562 | 0.497841 |
| | PC4 | Low | -0.03032 | -0.39171 | 1 |
| | PC5 | Low | -0.09317 | -1.00174 | 0.786614 |
| | PC6 | Low | -0.08794 | -0.7918 | 1 |
| | PC7 | Low | -0.13204 | -1.28715 | 0.289204 |
| Parkinson's disease | PC1 | High | 0.475532 | 1.523905 | 0.113516 |
| | PC2 | High | 0.084306 | 0.579395 | 1 |
| | PC3 | High | 0.084917 | 0.531798 | 1 |
| | PC4 | High | 0.412291 | 3.291441 | 0.00056 |
| | PC5 | High | 0.403644 | 1.445738 | 0.102365 |
| | PC6 | High | 0.241667 | 1.253787 | 0.415968 |
| | PC7 | High | 0.049625 | 0.220634 | 1 |
| | PC1 | Low | -0.09531 | -0.54773 | 1 |
| | PC2 | Low | -0.14351 | -1.29171 | 0.462954 |
| | PC3 | Low | -0.23413 | -1.17192 | 0.518192 |
| | PC4 | Low | -0.02665 | -0.26627 | 1 |
| | PC5 | Low | -0.18499 | -1.81221 | 0.157184 |
| | PC6 | Low | -0.03612 | -0.23002 | 1 |
| | PC7 | Low | -0.31817 | -2.78706 | 0.00056 |

**Supplementary Table 6**. Complete overview of risk estimates across all significant dimension–disease associations.

| Disease | PC | High/Low scores | Disease frequency in potentially high-risk group | Composite Score Threshold[1] | Disease frequency in potentially low-risk group | RR[2] total | RR female | RR male | PI[3] total |
|---|---|---|---|---|---|---|---|---|---|
| Cancer | PC1 | high | 456/9514 (4.792%) | > 0.232814 | 1570/ 61353 (2.558%) | 1.873 | 1.415 | 2.704 | 87.3 |
| | PC2 | low | 796/24668 (3.226%) | <-0.977425 | 409/48771 (2.889%) | 1.116 | 1.150 | 1.070 | 11.6 |
| | PC3 | high | 501/14463 (3.464%) | > 0.741493 | 1786/68099 (2.622%) | 1.321 | 1.088 | 1.708 | 32.1 |
| | PC4 | low | 607/19627 (3.092%) | < -0.889835 | 1276/44755 (2.851%) | 1.084 | 1.183 | 0 | 8.4 |
| | PC5 | high | 447/13554 (3.297%) | > 0.591762 | 1848/72528 (2.547%) | 1.294 | 1.107 | 2.572 | 29.4 |
| | PC6 | - | - | - | - | - | - | - | - |

| | | | | | | | | | |
|---|---|---|---|---|---|---|---|---|---|
| | PC7 | high | 799/25009 (3.194%) | > 0.683941 | 1662/62207 (2.671) | 1.195 | 1.180 | 1.234 | 19.5 |
| Diabetes | PC1 | high | 206/5099 (4.040%) | > 0.313200 | 1455/95804 (1.518%) | 2.661 | 2.669 | 2.668 | 166.1 |
| | PC2 | low | 507/26821 (1.890%) | < -0.962508 | 749/46157 (1.622%) | 1.165 | 1.211 | 1.119 | 16.5 |
| | PC3 | high | 347 /15054 (2.305%) | > 0.731554 | 1057/74491 (1.418%) | 1.625 | 1.622 | 1.631 | 62.5 |
| | PC4 | low | 427/17007 (2.510%) | < -0.931588 | 1173/83523 (1.404%) | 1.787 | 2.484 | 0 | 78.7 |
| | PC5 | high | 352/10837 (3.248%) | > 0.671768 | 1061/78064 (1.359%) | 2.389 | 2.217 | 4.409 | 138.9 |
| | PC6 | high | 566/24518(2.308%) | > 0.726202 | 960/63646 (1.508%) | 1.53 | 1.424 | 1.578 | 53 |
| | PC7 | high | 500/22998 (2.174%) | > 0.753079 | 1095/73350 (1.492%) | 1.457 | 1.428 | 1.636 | 45.7 |
| COPD | PC1 | high | 177/915 (19.344%) | > 0.481622 | 1018/14498 (7.021%) | 2.755 | 2.746 | 2.747 | 175.5 |
| | PC2 | - | - | - | - | - | - | - | - |
| | PC3 | low | 367 /3020 (12.152%) | < -0.878404 | 664 /10733 (6.186%) | 1.964 | 1.869 | 2.083 | 96.4 |
| | PC4 | high | 361/3781 (9.547%) | > 0.944709 | 688/9828 (7.000%) | 1.363 | 3.524 | 0.875 | 36.3 |
| | PC5 | high | 258/1772 (14.559%) | > 0.988681 | 931/13678 (6.806%) | 2.139 | 1.874 | 3.471 | 113.9 |
| | PC6 | high | 411/4246 (9.679%) | > 0.708331 | 771/10399 (7.414%) | 1.305 | 1.469 | 1.209 | 30.5 |
| | PC7 | high | 467/4806 (9.717%) | > 0.760298 | 788/11137 (7.075%) | 1.373 | 1.423 | 1.381 | 37.3 |
| Heart failure | PC1 | high | 145/949 (15.279%) | >0.365394 | 606/6941 (8.730%) | 1.75 | 2.182 | 1.414 | 75 |
| | PC2 | - | - | - | - | - | - | - | - |
| | PC3 | - | - | - | - | - | - | - | - |
| | PC4 | high | 487/4195 (11.609%) | > 0.650448 | 469/5485 (8.550%) | 1.357 | 2.082 | 0.789 | 35.7 |
| | PC5 | high | 158/1156 (13.667%) | > 0.878651 | 705 /8116 (8.686%) | 1.573 | 1.825 | 1.368 | 57.3 |
| | PC6 | high | 384/3494 (10.990%) | > 0.603445 | 612/6485 (9.437%) | 1.164 | 0.838 | 1.008 | 16.4 |
| | PC7 | low | 207/1763 (11.741%) | < -0.881934 | 655/6977 (9.387%) | 1.25 | 1.122 | 1.296 | 25 |
| Stroke | PC1 | high | 76/1069 (7.109%) | > 0.345871 | 350/10465 (3.344%) | 2.125 | 2.362 | 1.896 | 112.5 |
| | PC2 | - | - | - | - | - | - | - | - |
| | PC3 | - | - | - | - | - | - | - | - |
| | PC4 | high | 139/3246 (4.282%) | > 0.926495 | 261/7327 (3.562%) | 1.202 | 2.036 | 0.945 | 20.2 |
| | PC5 | - | - | - | - | - | - | - | - |
| | PC6 | - | - | - | - | - | - | - | - |
| | PC7 | - | - | - | - | - | - | - | - |

| | | | | | | | | | |
|---|---|---|---|---|---|---|---|---|---|
| Parkinson's disease | PC1 | - | - | - | - | - | - | - | - |
| | PC2 | - | - | - | - | - | - | - | - |
| | PC3 | - | - | - | - | - | - | - | - |
| | PC4 | high | 75/4293 (1.747%) | >0.579845 | 39/8766 (0.444%) | 3.934 | 6.920 | 0.903 | 293.4 |
| | PC5 | - | - | - | - | - | - | - | - |
| | PC6 | - | - | - | - | - | - | - | - |
| | PC7 | low | 53/3457 (1.533%) | < -0.53016 | 58/8870 (0.653%) | 2.347 | 1.935 | 2.231 | 134.7 |

1 Thresholds reflect the composite score value at which significant disease enrichment occurs, defining high-risk groups for each dimension.

2 RR: Relate Risk

3 PI: Percentage increase

### Supplementary Figures

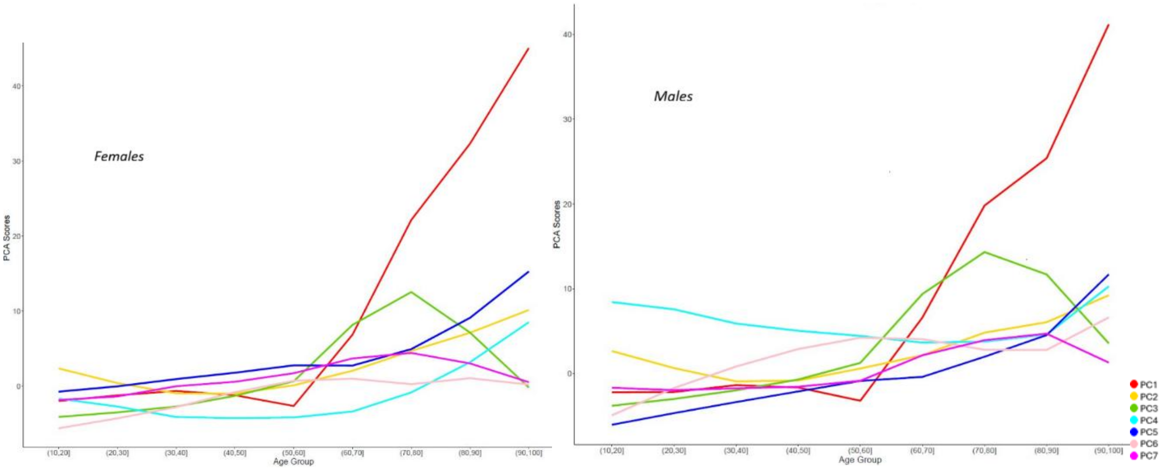


**Supplementary Figure 1.** Sex-specific differences in the seven aging-associated PCs.

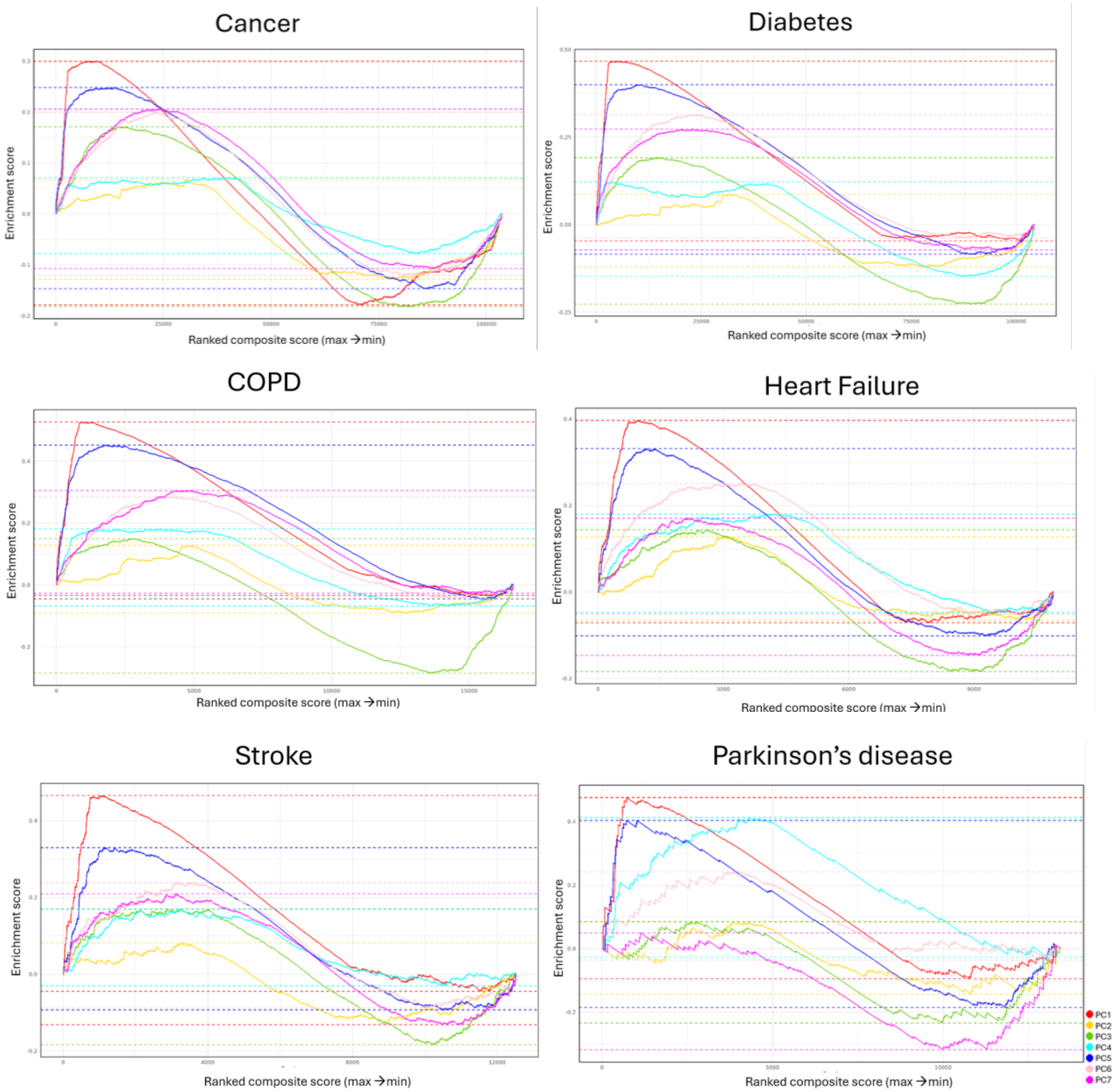


**Supplementary Figure 2.** Enrichment Scores (ES) of PC1-PC7 composite scores for cancer, diabetes, COPD, heart failure, stroke and Parkinson's disease incidence. The x-axis represents the ranked composite scores of participants (from maximum to minimum), and the y-axis indicates the ES. ES which is the maximum deviation from zero, indicates the magnitude and direction of enrichment. Positive and negative ES suggest overrepresentation at the high and low composite scores, respectively.

### Supplementary Notes

**Supplemnatry Note 1.** Participant and variable filtering

To construct a clean phenotypic dataset, participant- and variable-level filtering was performed based on data availability. Thresholds were selected following inspection of the distribution of missing data across the Lifelines dataset. Participants younger than 18 years were excluded. Variables were retained if data were available for more than 125,000 participants and if more than 25% of observations were non-missing. Application of these criteria resulted in a final analytical dataset comprising 152,241 participants and 1,177 phenotypic variables.
